# Biophysical modeling with variational autoencoders for bimodal, single-cell RNA sequencing data

**DOI:** 10.1101/2023.01.13.523995

**Authors:** Maria Carilli, Gennady Gorin, Yongin Choi, Tara Chari, Lior Pachter

## Abstract

We motivate and present *biVI*, which combines the variational autoencoder framework of *scVI* with biophysically motivated, bivariate models for nascent and mature RNA distributions. While previous approaches to integrate bimodal data via the variational autoencoder framework ignore the causal relationship between measurements, *biVI* models the biophysical processes that give rise to observations. We demonstrate through simulated benchmarking that *biVI* captures cell type structure in a low-dimensional space and accurately recapitulates parameter values and copy number distributions. On biological data, *biVI* provides a scalable route for identifying the biophysical mechanisms underlying gene expression. This analytical approach outlines a generalizable strategy for treating multimodal datasets generated by high-throughput, single-cell genomic assays.

## 1 Main

Advances in experimental methods for single-cell RNA sequencing (scRNA-seq) allow for the simultaneous quantification of multiple cellular species, such as nascent and mature transcriptomes [1,2], surface [3–5] and nuclear [6] proteomes, and chromatin accessibility [7,8]. While these rich datasets have the potential to enable unprecedented insight into cell type and state in development and disease, joint analyses of distinct modalities remain challenging. We show that principled biophysical “integration” of multimodal datasets can be achieved through parameterization of interpretable mechanistic models [9], scalable to measurements made for thousands of genes in tens of thousands of cells [10].

Recent approaches to integrate and reduce the dimensionality of multimodal single-cell genomics data have leveraged advances in machine learning [11–13]. For example, the popular tool *scVI* is a variational autoencoder (VAE) that uses neural networks to encode scRNA-seq counts to a low-dimensional representation. This is decoded by another neural network to a set of cell- and genespecific parameters for conditional likelihood distributions of observed counts. These distributions are chosen *post hoc* to be consistent with the discrete, over-dispersed nature of scRNA-seq counts, but can be derived from biophysical models (Section S1). Extensions of *scVI* to bimodal data have been attempted for protein [11] and chromatin measurements [14] by jointly encoding data modalities to a single latent space, then employing two decoding networks to produce parameters for *independent* conditional likelihoods specific to each datatype. Nascent and mature transcripts, available by realigning existing scRNA-seq reads [1, 2], could be similarly treated (Figure 1a). However, using independent conditional likelihoods for bimodal measurements derived from the same gene ignores the inherent causality between observations and has no biophysical basis: the generative model is merely part of a neural “black box” used to summarize data.

**Figure 1:**
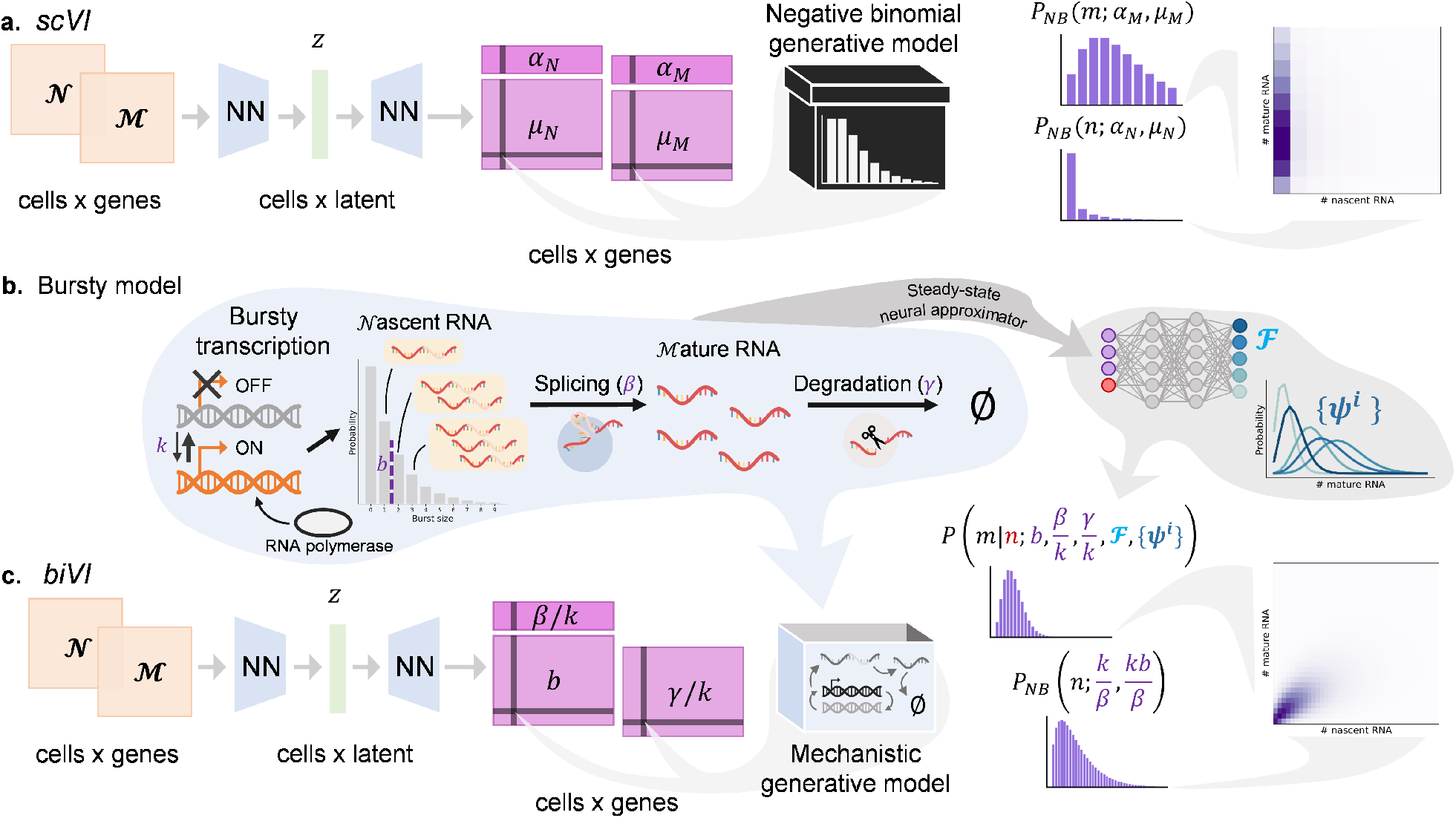
*biVI* reinterprets and extends *scVI* to infer biophysical parameters. **a**. *scVI* can take in concatenated nascent (𝒩) and mature (ℳ) RNA count matrices, encode each cell to a low-dimensional space *z*, and learn per-cell parameters *μ*_*N*_ and *μ*_*M*_ and per-gene parameters α _*N*_ and α_*M*_ for independent nascent and mature count distributions. This approach is not motivated by any specific biophysical model. **b**. A schematic of the telegraph model of transcription: a gene locus has the on rate *k*, the of rate *k*_*off*_, and the RNA polymerase binding rate *k*_RNAP_. Nascent RNA molecules are produced in geometrically distributed bursts with mean *b* = *k*_RNAP_*/k*_*off*_, which are spliced at a constant rate *β* and degraded at a constant rate *γ*. Although there is no closedform solution, this model’s steady-state distribution can be approximated by a pre-trained neural network ℱ and a set of basis functions {*ψ*^*i*^}. **c**. *biVI* can take in nascent and mature count matrices, produce a low-dimensional representation for each cell, and output per-cell parameters *b* and *γ*/*k*, as well as the per-gene parameters *β*/*k*, for a mechanistically motivated joint distribution of nascent and mature counts.

Nevertheless, good causal model candidates are available: for example, Figure 1b illustrates the extensively validated [15–17] bursty model of transcription. Nascent RNA molecules are produced in geometrically distributed bursts with mean *b* at constant rate *k* and spliced at rate*β* to produce mature molecules, which are degraded with constant rate*γ*. While the joint steady-state distribution induced by the bursty model is analytically intractable [18], we have previously shown that it can be approximated by a set of basis functions with neural-network learned weights [19].

We introduce *biVI*, a strategy that adapts *scVI* to work with well-characterized stochastic models of transcription. First, we propose several models, formalized by chemical master equations (CMEs), that could give rise to bivariate count distributions for nascent and mature transcripts. We then use the bivariate, CME-derived distribution as the conditional data likelihood distribution for nascent and mature counts (Figure 1c). The inferred conditional likelihood parameters thus have biophysical interpretations as part of a mechanistic model of transcriptional dynamics. Although we focus on the bursty model, *biVI* implements the closed-form constitutive and extrinsic noise models previously discussed in the literature [9, 20, 21] (derivations in Section S1 and diagrams in Figures S1 and S2).

After using simulations to show that *biVI* models, when compared to *scVI*, better recapitulate ground-truth distributions and achieve similar clustering of cell types’ latent representations (Figures S3, S4, S5, and Section S6.7), we applied *biVI* and *scVI* to experimental data [22] (Section 2.6) from mouse brain tissue [22]. As shown in Figure 2a-b, *biVI* recapitulates empirical distribution shapes better than *scVI* (Section 3) while allowing for interpretation of cell-specific parameters to determine *how* genes are regulated (Section S8). For example, in Figure 2c-d, we illustrate that the upregulation of markers *Foxp2* and *Rorb* can be ascribed to an increase in burst size. Wein Figure 2e, which shows the fraction of identified genes in each cell subclass that exhibited significant differences in burst size, relative degradation rate, or both (Section 2.8). Interesting trends across cell subclasses begin to emerge: neuronal cells appear to regulate gene expression via a mix of regulatory strategies, while non-neuronal cells seem to preferentially modulate burst size.

**Figure 2:**
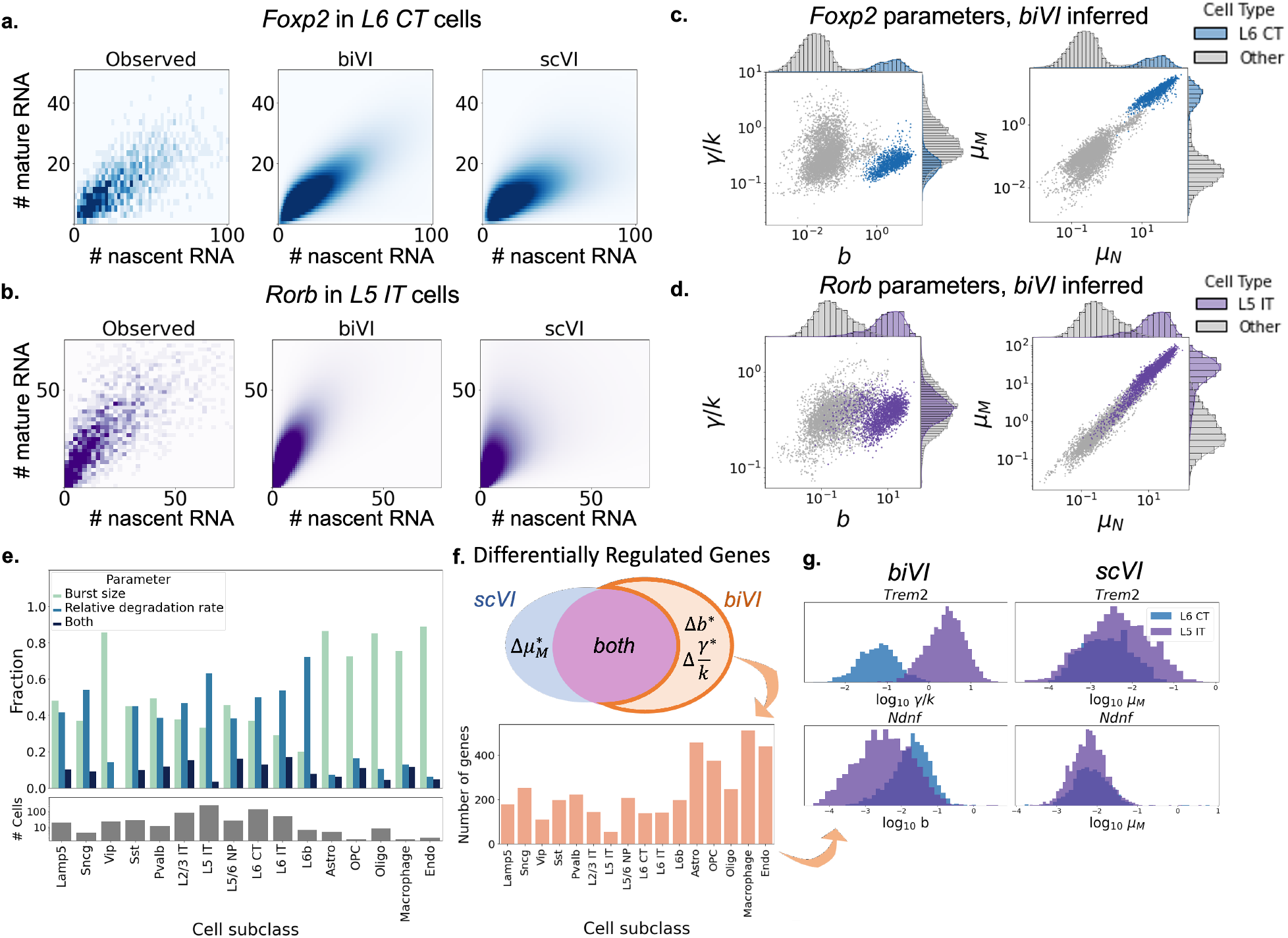
*biVI* successfully fits single-cell neuron data and suggests the biophysical basis for expression differences. **a**.-**b**. Observed, *scVI*, and *biVI* reconstructed distributions of *Foxp2*, a marker gene for L6 CT (layer 6 corticothalamic) cells, and *Rorb*, a marker gene for L5 IT (layer 5 intratelencephalic) cells, restricted to respective cell type. **c**.-**d**. Cell-specific parameters inferred for *Foxp2* and *Rorb* demonstrate identifiable differences in means and parameters in the marked cell types. **e**. Cell subclasses show different modulation patterns, with especially pronounced distinctions in non-neuronal cells (top: fractions of genes exhibiting differences in each parameter; bottom: number of cells in each subclass). **f**. *biVI* allows the identification of cells which exhibit differences in burst size or relative degradation rate, without necessarily demonstrating differences in mature mean expression. Hundreds of genes demonstrate this modulation behavior, with variation across cell subclasses. **g**. Histograms of *biVI* parameters and *scVI* mature means for two genes that exhibit parameter modulation without identifiable mature mean modulation. *Trem2* (top) shows differences in the degradation rate in *L5 IT* cells, whereas *Ndnf* (bottom) shows differences in burst size in *L6 CT* cells.

Finally, *biVI* can identify distributional differences which do not result in mean expression changes (Section 2.8, Figure 2g-h). For some cell subclasses, there were several hundred such genes, interesting targets for follow-up experimental investigation. For example, the gene *Ndnf*, which codes for the neuron derived neurotrophic factor NDNF, demonstrated a statistically significant difference in the *biVI* inferred burst size, but not *scVI* inferred mature mean, in the neuronal subclass L6 CT (Figure 2i, top row). NDNF promotes the growth, migration, and survival of neurons [23]; characterizing its regulatory patterns could help elucidate its role in neuronal maintenance. As another example, the relative degradation rate of the gene coding for the triggering receptor expressed on myeloid cells-2 (TREM2), variants of which are strongly associated with increased risk of Alzheimer’s disease [24], was found to be greater in the neuronal L5 IT subclass than in other subclasses (Figure 2i, bottom row). While known to be highly expressed in microglia [24], understanding its modulation in other cell subclasses could yield a better understanding of its cell type specific effects on the development of Alzheimer’s disease. Such mechanistic description provides a framework for characterizing the connection between a gene’s role and a cell’s regulatory strategies beyond a mere change in mean expression [25, 26].

We have demonstrated that bivariate distributions arising from mechanistic models can be used in variational autoencoders for principled integration of unspliced and spliced RNA-seq data. This improves model interpretability: conditional parameter estimates give insight into the mechanisms of gene regulation that result in differences in expression. While we impose biophysical constraints on species’ conditional joint distributions, orthogonal improvements in interpretability can be made by changing the decoder architecture. *biVI* models can be instantiated with single-layer linear decoders [27] to directly link latent variables with gene mean parameters via layer weights (Section S9 and Figure S9).

Relaxing assumptions and modeling more molecular modalities (e.g., protein counts and chromatin accessibility) are natural extensions. As single-cell technologies evolve to provide larger-scale, more precise measurements of biomolecules, we anticipate that our approach can be applied and extended for a more comprehensive picture of biophysical processes in living cells.

## 2 Methods

In order to extend the *scVI* method to work with multimodal molecule count data in a way that is coherent with biology, we define bivariate likelihood functions that (i) encode a specific, precedented mechanistic model of transcriptional regulation and (ii) are admissible under the assumptions made in the standard *scVI* pipeline. On a high level, our method entails the following steps:

1. Choose one of the *scVI* univariate generative models (Section 2.2), including the functional form of its likelihood and any assumptions about its distributional parameters.
2. Identify a one-species chemical master equation (CME) that produces this distribution as its steady state, and translate assumptions about distributional parameters into assumptions about the biophysical quantities that parameterize the CME (Section 2.3). The one-species system and its assumptions will typically not be uniquely determined.
3. Identify a two-species CME and derive assumptions about parameter values consistent with the one-species system (Section 2.3). There will typically be multiple ways to preserve the assumptions but only a single CME.
4. Modify the autoencoder architecture to output the variables that parameterize the CME solution under the foregoing assumptions, and use this solution as the generative model (Section 2.5).

### 2.1 Statistical preliminaries

We use the standard parameterization of the Poisson distribution:

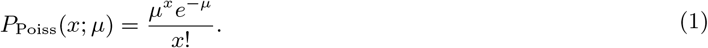

We use the shape-mean parameterization of the univariate negative binomial distribution:

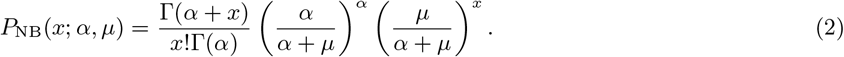

We use mean parameterization of the geometric distribution on ℕ_0_:

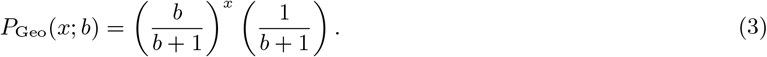

### 2.2 *scVI* models

A brief summary of the generative process of the standard, univariate *scVI* pipeline is useful to contextualize the options and constraints of the bivariate model. In the Bayesian model, each cell has some posterior probability *p_c_*(*z_c_*) over a low-dimensional space and can be represented as a sample *z_c_* from that posterior. *scVI* uses the “decoder” neural network to map from realizations *z_c_* to quantities *ρ*_*cg*_, which describe the compositional abundance of gene *g* in cell *c* as a function of *z_c_*, such thatΣ*g ρ_cg_* = 1. Furthermore, a cell-specific “size factor” *ℓ*_*c*_ is sampled from a lognormal distribution parameterized by either fit or plug-in estimates of mean and variance such that the mean expression of a gene in a given cell is *μ*_*cg*_ = *ρ*_*cg*_*ℓ*_*c*_.

The univariate workflow provides the options of three discrete generative models: Poisson with mean *μ*_*cg*_, negative binomial with mean *μ*_*cg*_ and gene-specific dispersion parameter *α*_*g*_, and zero-inflated negative binomial, with an additional Bernoulli mixture parameter. We report the master equation models consistent with the first two generative laws below, and discuss a potential basis for and reservations about the zero-inflated model in Section S1.4.

Due to the intractability of the posterior probability *p*_*c*_(*z*_*c*_), *scVI* uses variational inference to infer an approximate posterior *q*_*z*_ (*z*_*c*_), which is in form a multivariate Gaussian. Models are trained via stochastic optimization of the Evidence Lower Bound, or ELBO, which minimizes the Kullback-Leibler divergence between the approximate posterior and a prior and maximizes the expectation value of conditional likelihood over the approximate posterior. The Gaussian form of the approximate posterior makes possible a reparameterization trick to calculate gradients of the ELBO over expectation estimates made by Monte Carlo sampling from the approximate posterior. Further, the encoding network amortizes inference by learning a map between data to parameters of the approximate posterior [12, 28].

### 2.3 Master equation models

The one-species CMEs encode reaction schema of the following type:

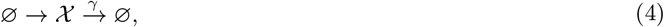

where 𝒳 is a generic transcript species used to instantiate a univariate *scVI* generative model, *γ* is the transcript’s Markovian degradation rate, and the specific dynamics of the transcription process (first arrow) are deliberately left unspecified for now. Such systems induce univariate probability laws of the form *P* (*x*).

The two-species CMEs encode reaction schema of the following type:

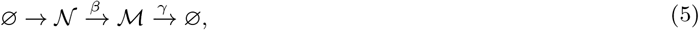

where 𝒩 denotes a *n*ascent species, ℳ denotes a *m*ature species, and *β* denotes the nascent species’ Markovian conversion rate. Such systems induce bivariate probability laws of the form *P* (*n, m*). We typically identify the nascent species with unspliced transcripts and the mature species with spliced transcripts. We use the nascent/mature nomenclature to simplify notation and emphasize that this identification is natural for scRNA-seq data, but not mandatory in general.

Formalizing a model in terms of the CME requires specifying the precise mechanistic meaning of *ρ*_*cg*_ and *ℓ*_*c*_. Previous reports equivocate regarding the latter [11], appealing either to cell-wide effects on the biology (in the spirit of [20, 21]) or technical variability in the sequencing process (in the spirit of [29]). For completeness, we treat both cases.

Below, we present the theoretical results, including the biophysical models, the functional forms of bivariate distributions consistent with the standard *scVI* models, and the consequences of introducing further assumptions. The full derivations are given in Section S1

#### 2.3.1 *Constitutive*: The Poisson model and its mechanistic basis

The Poisson generative model can be recapitulated by the following schema:

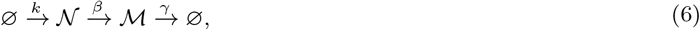

where *k* is a constant transcription rate. This process converges to the bivariate Poisson stationary distribution, with the following likelihood:

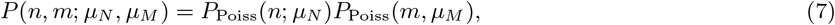

where *μ*_*N*_ = *k/ β* and *μ*_*M*_ = *k/ γ*. If we suppose each gene’s *β* and *1* are constant across cell types, the likelihoods involve a single compositional parameter *ρ*_*cg*_, such that

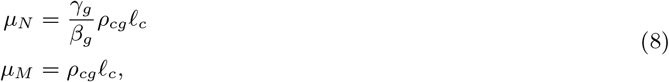

where *γ*_*g*_/*β*_*g*_ ∈ ℝ^+^ is a gene-specific parameter that can be fit or näively estimated by the ratio of the unspliced and spliced averages. On the other hand, if the downstream processes’ kinetics can also change between cell types, we must use two compositional parameters:

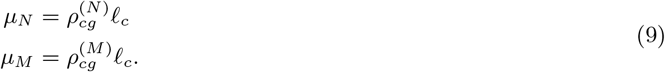

We refer to this model as “Poisson,” reflecting its functional form, or “constitutive,” reflecting its biophysical basis.

#### 2.3.2 *Extrinsic*: The negative binomial model and a possible mixture basis

The negative binomial generative model can be recapitulated by the following schema:

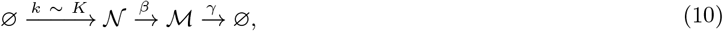

where *k* is the transcription rate, a realization of *K*, a gamma random variable with shape *λ*, scale *η*, and mean ⟨*K*⟩ = *λη*. This process converges to the bivariate negative binomial (BVNB) stationary distribution, with the following likelihood:

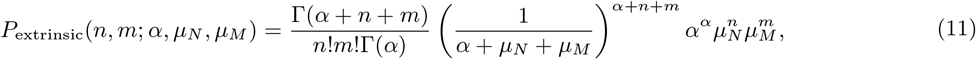

where *μ*_*N*_ =⟨*K*⟩/*β* and *μ*_*M*_ = ⟨*K*⟩/*γ*. If we suppose that cell type differences only involve changes in the transcription rate scaling factor *η*, with constant *α*,/*β*, and *γ*, the likelihoods involve a single compositional parameter *ρ*_*cg*_. The mean parameters are identical to Equation 8, with an analogous parameter *γ*_*g*_/*β*_*g*_, as well as a gene-specific shape parameter *α*_*g*_. On the other hand, if the downstream processes’ kinetics can also change between cell types, we must use two compositional parameters, as in Equation 9.

We refer to this model as “extrinsic” to reflect its biophysical basis in extrinsically stochastic rates of transcriptional initiation.

#### 2.3.3 *Bursty*: The negative binomial model and a possible bursty basis

The negative binomial generative model may be recapitulated by the alternative schema [18]:

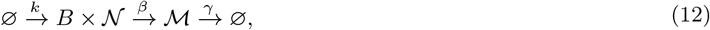

where *k* is the burst frequency and *B* is a geometric random variable with mean *b* (Equation 3). This system converges to the following stationary distribution:

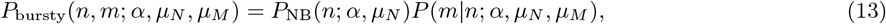

where *μ*_*N*_ = *kb/β, μ*_*M*_ = *kb/γ*, andα is arbitrarily set to *k/β* for simplicity.

Although the nascent marginal is known to be negative binomial, the joint *P* (*n, m*) and conditional *P* (*m*|*n*) distributions are not available in closed form. For a given set of parameters, the joint distribution can be approximated over a finite microstate domain *n, m ∈* [0, ß_*N*_ − 1] × [0, ß_*M*_ − 1], with total *s*tate *s*pace size ß_*N*_ ß_*M*_. This approach is occasionally useful, if intensive, for evaluating the likelihoods of many independent and identically distributed samples. The numerical procedure entails using quadrature to calculate values of the generating function on the complex unit sphere, then performing a Fourier inversion to obtain a probability distribution [18]. However, this strategy is inefficient in the variational autoencoder framework, where each observation is associated with a distinct set of parameters. Furthermore, it is incompatible with automatic differentiation.

In [19], we demonstrated that the numerical approach can be simplified by approximating *P* (*m*|*n*) with a learned mixture of negative binomial distributions: the weights are given by the outputs of a neural network, whereas the negative binomial bases are constructed analytically. The neural network is trained on the outputs of the generating function procedure. Although the generative model does not have a simple closed-form expression, it is represented by a partially neural, pre-trained function that is *a priori* compatible with the VAE.

If we suppose cell type differences only involve changes in the burst size *b*, with constant *k,/β* and *γ*, we use Equation 13 to evaluate likelihoods. These likelihoods involve a single compositional parameter *ρ*_*cg*_, with mean parameters identical to Equation 8, with an analogous parameter *γ*_*g*_/*β*_*g*_, as well as a gene-specific shape parameter *α*_*g*_. On the other hand, if kinetics of the degradation process can also change between cell types, we must use two compositional parameters, as in Equation 9. There is no admissible way to allow modulation in the burst frequency.

We refer to this model as “bursty,” reflecting its biophysical basis.

### 2.4 *biVI* bursty generative model

Following the notation of *scVI* [28], *biVI* ‘s generative process for the bursty hypothesis models expression values of *x*_*cn*_ and *x*_*cm*_ of nascent and mature counts, respectively, in cell *c* as:

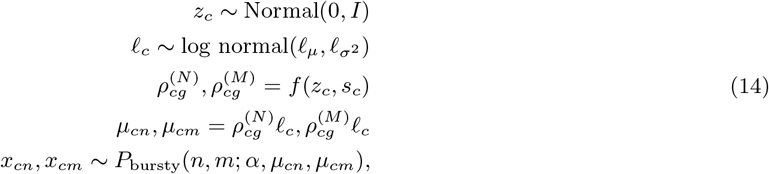

with a standard, multivariate normal prior on the latent space *z* vector. Here, *ℓ*_*μ*_, 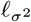 are by default observed mean and variance in log-sequencing depth (‘log-library size’ in *scVI*) across a cell’s batch, although they can be learned. Further, as in *scVI, f* is neural network that produces fraction of sequencing depth parameters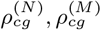for nascent and mature counts. The sum of nascent and mature fractions is constrained to be 1 over a cell *c* by a softmax applied to the network output: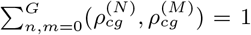 where *G* is the number of genes. *α* ∈ ℝ ^*G*^ is a network parameter jointly optimized across all cells during the variational inference procedure. To recover biophysical parameters, *α* is arbitrarily set to *k/β*. Burst size *b* and relative degradation rate *k/ γ* can be recovered according to the following conversions:

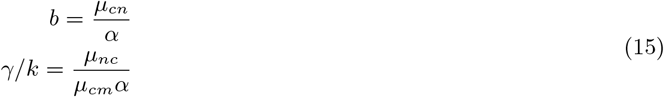

We further set *k* = 1 with no loss of generality at steady-state. Generative processes for constitutive and extrinsic noise models are discussed in Sections S2 and S3.

### 2.5 *biVI* modifications to *scVI*

Our code is built upon *scVI* version 0.18.0 [30]; the following outlines the modifications we made for *biVI*. The *scVI* framework already supports the constitutive model. By setting conditional likelihood to “poisson,” no modification of *scVI* architecture is necessary. The conditional data likelihood distribution is the product of two Poisson distributions (Equation 7). Explicitly, unspliced and spliced count matrices can be concatenated along the cell axis to produce a matrix of shape *C* by 2*G*, where *C* is the number of cells and *G* the number of genes. *scVI* will then produce 2*G* Poisson mean parameters for the two Poisson distributions of Equation 7.

For the extrinsic and bursty models, mean parameters for nascent and mature counts, *μ*_*N*_ and *μ*_*M*_, and a single shape parameter *ff* are necessary. The default *scVI* architecture returns two independent parameters for nascent and mature counts of the same gene. *biVI* thus modifies the *scVI* architecture to update vectors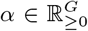 rather than 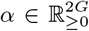 where *G* is the number of genes. For the extrinsic model, the conditional data likelihood distribution is set to the extrinsic likelihood *P*_extrinsic_(*n, m*; *α, μ*_*N*_, *μ*_*M*_) (Equation 11). For the bursty model, the conditional data likelihood distribution is set to the bursty likelihood *P*_bursty_(*n, m*; *α,μ*_*N*_, *μ*_*M*_) (Equation 13). These models also intake concatenated unspliced and spliced matrices of shape *C* by 2*G*

### 2.6 Preprocessing Allen data

Raw 10x v3 single-cell data were originally generated by the Allen Institute for Brain Science [22]. The raw reads in FASTQ format [31] and cluster metadata [32] were obtained from the NeMO Archive. We selected mouse library B08 (donor ID 457911) for analysis.

To obtain spliced and unspliced counts, we first obtained the pre-built mm10 mouse genome released by 10x Genomics (https://support.10xgenomics.com/single-cell-gene-expression/software/downloads/latest, version 2020-A). We used *kallisto*|*bustools* 0.26.0 [2] to build an intronic/exonic reference (`kb ref` with the option `--lamanno`). Next, we pseudoaligned the reads to this reference (`kb count` with the option `--lamanno`) to produce unspliced and spliced count matrices. We used the outputs produced by the standard *bustools* filter. This filter was relatively permissive: all (8,424) barcodes given cell type annotations in the Allen metadata were present in the output count matrix (10,975 barcodes).

Based on previous clustering results, we selected cells that were given cell type annotations, and omitted “low quality” or “doublet” barcodes [22], for a total of 6,418 cells. Although any choice to retain or omit cells from analysis is arbitrary, our work models the generating process that produced cells’ nascent and mature counts by presupposing each barcode corresponds to a single cell. Therefore, we propose that cells identified as low-quality (empty cells) or as doublets (two cells measured in one observation) [22] have a fundamentally different data-generating process than individual single cells, and therefore remove them before fitting VAE models. However, we stress that the stochastic nature of transcription and sequencing, the intrinsic uncertainties associated with read alignment, and the numerical compromises made in clustering large datasets mean that previous annotations are not “perfect,” merely a reasonable starting point for comparing alternative methods.

We used Scanpy [33] to restrict our analysis to the most variable genes, which presumably reflect the cell type signatures of interest. The spliced count matrix for the 6,418 retained cells was normalized to sum to 10,000 counts per cell, then transformed with log1p. The top 2,000 most highly variable genes were identified using`scanpy.pp.highly_variable_genes` on spliced matrices with minimum mean of 0.0125, maximum mean of 3, and minimum dispersion of 0.5 [33]. Spliced and unspliced matrices were subset to include only the 2,000 identified highly variable genes, then concatenated along the cell axis in the order unspliced, spliced to produce a count matrix of size 6,418 by 4,000.

### 2.7 Fitting Allen data

We applied *biVI* with the three generative models (bursty, constitutive, and extrinsic) and *scVI* with negative binomial likelihoods to the concatenated unspliced and spliced count matrix obtained by the filtering procedures outlined above. We made the key assumption that unspliced and spliced counts could be treated as the nascent and mature species of the bursty generative model (see discussion in Section S5). 4,622 cells were used for training with 513 validation cells, and 1,283 cells were held out for testing performance. All models were trained for 400 epochs with a learning rate of 0.001. Encoders and decoder consisted of 3 layers of 128 nodes, and each model employed a latent dimension of 10

### 2.8 Bayes factor hypothesis testing for differential expression

After fitting the VAE models, we sought to identify meaningful statistical differences that distinguish cell types. We excluded cell subclasses “L6 IT Car3,” “L5 ET,” “VLMC,” and “SMC” from this analysis, as they contained fewer than ten annotated cells and may require more sophisticated statistical models to account for small sample sizes. The following analysis thus considers 6,398 cells in 16 unique subclasses. We only computed differential expression metrics under the bursty model.

Differential parameter values were tested for each assigned subclass label (as annotated in [22]) versus all others using a Bayes factor hypothesis test following [11]. We reproduce Equations (18) - (21) of [11] below for clarity.

Estimating differential values of any parameter *θ*^*g*^ of gene *g* in cells *a* and *b* can be done according to the following Bayesian framework. First, as in Equation (18) of [11], the log fold change (LFC) of *θ*^*g*^ between two cells *a* and *b* can be calculated as follows:

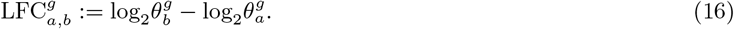

Then, as in Equation (19) of [11], the probability that the magnitude of the LFC is greater than some effect threshold *T* can be found by evaluating 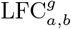over the posterior distributions of each cell:

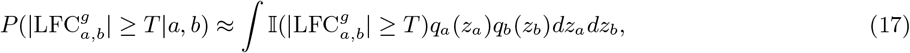

where, in practice, the integral is approximated with many Monte Carlo samples from the two cells’ posteriors. Two hypotheses are tested: *H*_1_, or that the magnitude of the LFC is greater than or equal to threshold *T*, and *H*_0_, or the null hypothesis that the magnitude of the LFC is less than *T*. A Bayes factor for gene *g* between cells *a* and *b*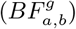 is calculated to compare the two hypotheses, as in Equation (20) of [11]:

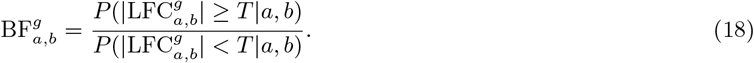

Extending this to test differential expression between two groups of cells *A* and *B* amounts to “aggregating the posterior,” as in Equation (21) of [11], or evaluating the same 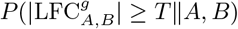 over

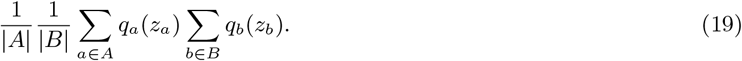

In other words, a random sample *z*_*a*_ can be be taken from the approximate posterior of any cell belonging to group *A* and decoded to produce parameter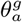 likewise a random sample *z*_*b*_ can be taken from the approximate posterior of any cell belonging to group *B* and decoded to produce parameter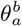. The LFC between the two parameters can then be calculated. Repeating this for many Monte Carlo samples over the aggregate posteriors allows estimation of the Bayes factor between two groups.

For the results shown in Figure 1, we used cutoffs of *T* ≥ 1.0, or a magnitude LFC of ≥ 2, and a Bayes factor threshold of 1.5. The Bayes factors were calculated on normalized burst size and means for *biVI*, i.e., the fractional inferred burst size or inferred means (before scaling by sampled sequencing depth for that cell), and normalized means for *scVI*. This controlled for differences in parameters due to sequencing depth that were not biologically meaningful. Relative degradation rate*γ/k* is independent of sequencing depth: hypothesis tests were performed directly on inferred relative degradation rates. While batch identity can also be integrated over to compare groups of cells from different batches, our analysis did not require this as all cells were from the same batch.

## 3 Reconstructing gene distributions

Let *θ*_*kg*_ be mechanistic model parameters for gene *g* in cell type *k*. While parameters for a given gene are identical across all cells in a specific cell type, *biVI* and *scVI* infer unique parameters for every cell and gene: *θ*_*cg*_, where *c* indexes over cells and *g* indexes over genes. To reconstruct distributions for a given gene in a specific cell type *k*, we sample once from the posterior distribution *q*_*c*_(*z*) of each cell *c*∈*k* to obtain point-estimates of conditional parameter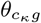 where conditional refers to a single sampling from a cell’s posterior, or a particular realization of *z*_*c*_. We then average over the cell-specific conditional probabilities for the gene to produce a cell type marginal distribution:

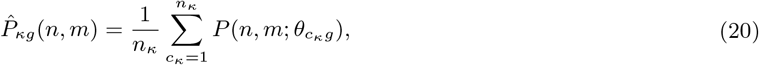

where *n*_*k*_ is the total number of cells in cell type *k*, and *c*_*k*_ indexes over all cells in that cell type. This identity follows immediately from defining the cell type’s distribution as the mixture of the distributions of its constituent cells. In the case of *biVI*, we plug in Equation 7, 11, or 13 for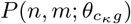. In the case of *scVI*, we use a product of two independent negative binomial laws:

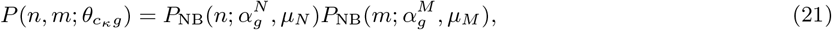

where *μ*_*N*_ and *μ*_*M*_ are cell- and gene-specific, whereas *α*^*N*^ and *α* ^*M*^ are fit separately and take on different values (Section 2.5). For simplicity, this comparison omits uncertainty associated with *θ*_*cg*_, which is formally inherited from the uncertainty in the latent representation *z* for each cell *c*.

Thus, Equation 21 is an approximation to the posterior predictive distribution, or marginal distribution of data given the approximated posterior, if we assume Monte Carlo sampling from the approximate posterior distributions of cells within that cell type as a reasonable proxy for sampling from the cell type’s posterior distribution. The posterior predictive, or marginal, distribution is:

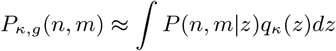

where *q*(*z*) is the approximate posterior. We further note that conditional data likelihood and the marginal distribution are *not* necessarily of the same form (for example, if the conditional data likelihood distribution is negative binomial, the marginal distribution of genes is not necessarily negative binomial).

## 4 Data availability

Simulated datasets, simulated parameters used to generate them, and Allen dataset B08 and its associated metadata are available in the Zenodo package 7497222. All analysis scripts and notebooks are available at https://github.com/pachterlab/CGCCP_2023. The repository also contains a Google Colaboratory demonstration notebook applying the methods to a small human blood cell dataset.

## 5 Acknowledgments

M.C., G.G., T.C., and L.P. were partially funded by NIH 5UM1HG012077-02 and NIH U19MH114830. Y.C. was partially funded by T32 GM007377. G.G. thanks Drs. Ido Golding and Heng Xu for the inspiration leading to the explanatory model for the zero-inflated negative binomial distribution in Section S1.4. The RNA illustrations used in Figures 1, S1, and S2 were derived from the DNA Twemoji by Twitter, Inc., used under the CC-BY 4.0 license. We thank the Caltech Bioinformatics Resource Center for GPU resources that helped in performing the analyses.

