## Supplemental material for "Biophysical modeling with variational autoencoders for bimodal, single-cell RNA sequencing data"

May 2, 2023

### S1 Derivations

As discussed in Section 2.3, our models encode a series of single-molecule stochastic processes.

#### S1.1 The Poisson model and its constitutive extensions

The simplest standard *scVI* generative model is Poisson (Equation 1), where $\mu = \mu_{cg}$ is cell- and gene-specific. Furthermore, $\mu_{cg} = \rho_{cg}(z)\ell_c$ encodes dependence on the latent encoding $z$ of cell $c$ through the compositional gene abundance $\rho_{cg}$ (such that $\sum_g \rho_{cg} = 1$), as well as the genome-wide "cell size" scaling factor $\ell_c$.

The following model produces a Poisson stationary distribution of $\mathcal{X}$ [34]:

$$\varnothing \xrightarrow{k} \mathcal{X} \xrightarrow{\gamma} \varnothing. \tag{22}$$

This is the standard model of constitutive, or unregulated, transcription [35] at a single gene locus, with mean $\mu = k/\gamma$.

##### S1.1.1 "Cell size" as biological variation

The parameter $k$ is the transcription rate. In physical terms, it typically represents the rate of RNA polymerase (RNAP) recruitment at the gene locus. The rate of recruitment may be modeled by a simple mass action model:

$$k_{cg} = k_{cg,init}[\text{RNAP}]_c. \tag{23}$$

Thus, the concentration of RNAP is cell-specific, and controls the transcription rate for all genes. On the other hand, the initiation rate differs between cells (e.g., due to different levels of chromatin accessibility) as well as different genes (e.g., due to differences in binding kinetics between promoter motifs). We may choose the units of concentration to yield $[\text{RNAP}]_c \sim \mathcal{O}(1)$. This formulation implies:

$$\mu_{cg} = \frac{k_{cg}}{\gamma_g} = \frac{k_{cg,init}}{\gamma_g}[\text{RNAP}]_c = \frac{k_{z,init}}{\gamma_g}[\text{RNAP}]_c, \tag{24}$$

where we assume $\gamma$ is constant across different cells.

The first factor encodes transcriptional modulation due to the specific gene identity and location in latent space, whereas the second factor encodes transcriptional modulation due to "cell size" effects. If we include an arbitrary, but large universal scaling factor $C$ (on the order of $10^4$), we obtain

$$\mu_{cg} = \frac{k_{z,init}}{C\gamma_g}C[\text{RNAP}]_c, \tag{25}$$

and can *define* the *scVI* factors as:

$$\begin{aligned} \rho_{cg} &:= \frac{k_{z,init}}{C\gamma_g}, \\ \ell_c &:= C[\text{RNAP}]_c, \end{aligned} \tag{26}$$

which accords with the premise that the values of $\rho_{cg}$ are near zero, whereas those of $\ell_c$ are quite large and comparable to the cells' total UMI counts.

Next, we write down the two-species system:

$$\varnothing \xrightarrow{k} \mathcal{N} \xrightarrow{\beta} \mathcal{M} \xrightarrow{\gamma} \varnothing, \tag{27}$$

which induces the bivariate Poisson stationary distribution:

$$P(n, m; \mu_N, \mu_M) = P_{\text{Poiss}}(n; \mu_N) P_{\text{Poiss}}(m; \mu_M), \tag{28}$$

where $\mu_N = k/\beta$ and $\mu_M = k/\gamma$. We maintain the assumptions of $z$ and "cell size" dependence through $k$, and impose the assumption that both $\beta$ and $\gamma$ are gene-specific and constant. This yields:

$$\begin{aligned} \mu_{cg}^{(N)} &= \frac{k_{z,init}}{C\beta_g} C[\text{RNAP}]_c = \frac{\gamma_g}{\beta_g} \rho_{cg} \ell_c, \\ \mu_{cg}^{(M)} &= \frac{k_{z,init}}{C\gamma_g} C[\text{RNAP}]_c = \rho_{cg} \ell_c, \end{aligned} \tag{29}$$

where $\gamma_g/\beta_g$ may be fit or estimated from the plug-in method of moments estimate $\overline{N}_g/\overline{M}_g$. On the other hand, if we suppose that $\gamma$ and $\beta$ can differ between cell types, we are forced to define two distinct $\rho$ parameters:

$$\begin{aligned} \mu_{cg}^{(N)} &= \rho_{cg}^{(N)} \ell_c \text{ s.t. } \rho_{cg}^{(N)} := \frac{k_{z,init}}{C\beta_z}, \\ \mu_{cg}^{(M)} &= \rho_{cg}^{(M)} \ell_c \text{ s.t. } \rho_{cg}^{(M)} := \frac{k_{z,init}}{C\gamma_z}. \end{aligned} \tag{30}$$

In sum, if we assume each gene's splicing and degradation rates are constant across all cells, we use Equation 28 to evaluate likelihoods, with $\mu_N = \frac{\gamma_g}{\beta_g} \rho_{cg} \ell_c$ and $\mu_M = \rho_{cg} \ell_c$. On the other hand, if we relax this assumption, we use the same equation, with $\mu_N = \rho_{cg}^{(N)} \ell_c$ and $\mu_M = \rho_{cg}^{(M)} \ell_c$.

##### S1.1.2 "Cell size" as technical variation

We can recapitulate the same quantitative model by describing $\ell_c$ as reflective of a technical phenomenon. First, we replace $\mathcal{X}$ with $\mathcal{X}_b$, the *biological* molecules. Next, we impose the hypothesis that each molecule of $\mathcal{X}_b$ has probability $p$ of actually being captured, amplified, and sequenced during the experiment. The sequencing process induces a Poisson stationary distribution of observed counts $\mathcal{X}$, with mean $\mu = kp/\gamma$. This model can be obtained from a model of conversion $\mathcal{X}_b \to \mathcal{X}$ or in the low-rate limit of a model of catalytic production $\mathcal{X}_b \to \mathcal{X}_b + \mathcal{X}$ [36].

Therefore, we can parameterize a simple generative model of imperfect capture:

$$\mu_{cg} = \frac{k_{cg}}{\gamma_{cg}} p_c = \mu_{cg}^b p_c, \tag{31}$$

i.e., downsampling amounts to rescaling the expectation of the *b*iological quantity. The assumption that $p_c$ does not vary between genes reflects a specific class of hypotheses, e.g., poly(A) tails being chemically similar in a sequencing experiment. This model may also reflect more generic descriptions of variation, as the factors of $\mu_{cg}$ are not independently identifiable. For example, if $p_{cg} = p_c\xi_g$ for some coefficient $\xi_g$ of moderate magnitude, we merely find that $\mu^b_{cg}$ is rescaled by $\xi_g$.

If we introduce the scaling factor $C$, we obtain

$$\mu_{cg} = \frac{\mu^b_{cg}}{C} C p_c. \tag{32}$$

Much as before, if $C$ is relatively large, we may define the variables as:

$$\begin{aligned} \rho_{cg} &:= \frac{\mu^b_{cg}}{C}, \\ \ell_c &:= C p_c. \end{aligned} \tag{33}$$

In the two-species case (Equation 27), the same identities hold [37]. The bivariate law is given by Equation 28 with $\mu_N = \mu^b_N p_c$ and $\mu_M = \mu^b_M p_c$. As above, the assumption of a common $p_c$ is not mandatory, but the functional form can be derived by proposing $p^{(N)}_{cg} = \xi^{(N)}_{cg} p_c$ and $p^{(M)}_{cg} = \xi^{(M)}_{cg} p_c$, followed by rescaling the biological expectations. Thus, if $\beta$ and $\gamma$ can differ, we find:

$$\begin{aligned} \mu^{(N)}_{cg} &= \frac{k_z}{\beta_z}\ell_c = \rho^{(N)}_{cg}\ell_c, \\ \mu^{(M)}_{cg} &= \frac{k_z}{\gamma_z}\ell_c = \rho^{(M)}_{cg}\ell_c. \end{aligned} \tag{34}$$

This distinct set of assumptions recapitulates the functional form in the previous section. To obtain this solution, we used a convenient property by which a Bernoulli-subsampled Poisson distribution remains Poisson.

#### S1.2 The negative binomial model and its mixture extensions

The *scVI* negative binomial generative model is based on (Equation 2), where $\mu = \mu_{cg}$ is defined as above, whereas $\alpha = \alpha_g$ is the shape or "inverse dispersion" parameter, presumed to be gene-specific.

The following model produces a negative binomial stationary distribution of $\mathcal{X}$:

$$\varnothing \xrightarrow{k \sim K} \mathcal{X} \xrightarrow{\gamma} \varnothing, \tag{35}$$

where the transcription rate $k$ is drawn from a gamma distribution with shape $\alpha$, scale $\eta$, and mean $\langle K \rangle = \alpha\eta$. This is the mixture extension of the constitutive model, which encodes variability in the transcription rate [20, 21], potentially due to an unspecified, but slow, stochastic process [9]; the mean of this distribution is $\langle K \rangle / \gamma$.

##### S1.2.1 "Cell size" as biological variation

To include dependence on $z$ and the "cell size," while maintaining the assumption of a constant shape parameter, we can generalize the model in Section S1.1.1. First, we note that $\eta$, the gene-specific scale parameter that governs the frequency of transcriptional events, must be proportional to the concentration term [RNAP] [9]. By choosing the appropriate units and defining $\eta_{cg} =$

$k_{z,init}[\text{RNAP}]_c$, we obtain

$$\begin{aligned} \mu_{cg} &= \frac{\alpha_g k_{z,init}}{C\gamma_g} C[\text{RNAP}]_c, \\ \rho_{cg} &:= \frac{\alpha_g k_{z,init}}{C\gamma_g}, \\ \ell_c &:= C[\text{RNAP}]_c, \end{aligned} \tag{36}$$

which is consistent with the Poisson result. Next, we write down the two-species system:

$$\varnothing \xrightarrow{k \sim K} \mathcal{N} \xrightarrow{\beta} \mathcal{M} \xrightarrow{\gamma} \varnothing, \tag{37}$$

which induces the bivariate negative binomial stationary distribution [38]:

$$P(n, m; \alpha, \mu_N, \mu_M) = \frac{\Gamma(\alpha + n + m)}{n!m!\Gamma(\alpha)} \left(\frac{\alpha}{\alpha + \mu_N + \mu_M}\right)^{\alpha} \left(\frac{\mu_N}{\alpha + \mu_N + \mu_M}\right)^{n} \left(\frac{\mu_M}{\alpha + \mu_N + \mu_M}\right)^{m}. \tag{38}$$

As in the generic case, $\mu_N = \langle K \rangle / \beta$ and $\mu_M = \langle K \rangle / \gamma$. We maintain the assumptions of $z$ and "cell size" dependence through $k$, and impose the assumption that both $\beta$ and $\gamma$ are gene-specific and constant. This yields:

$$\begin{aligned} \mu_{cg}^{(N)} &= \frac{\alpha_g k_{z,init}}{C\beta_g} C[\text{RNAP}]_c = \frac{\gamma_g}{\beta_g} \rho_{cg} \ell_c, \\ \mu_{cg}^{(M)} &= \frac{\alpha_g k_{z,init}}{C\gamma_g} C[\text{RNAP}]_c = \rho_{cg} \ell_c, \end{aligned} \tag{39}$$

where $\gamma_g/\beta_g$ may be fit or estimated from the plug-in method of moments estimate $\overline{N}_g/\overline{M}_g$. As before, if we suppose that $\gamma$ and $\beta$ can differ between cell types, we are forced to define two distinct $\rho$ parameters:

$$\begin{aligned} \mu_{cg}^{(N)} &= \rho_{cg}^{(N)} \ell_c \text{ s.t. } \rho_{cg}^{(N)} := \frac{\alpha_g k_{z,init}}{C\beta_g}, \\ \mu_{cg}^{(M)} &= \rho_{cg}^{(M)} \ell_c \text{ s.t. } \rho_{cg}^{(M)} := \frac{\alpha_g k_{z,init}}{C\gamma_g}. \end{aligned} \tag{40}$$

In sum, if we assume each gene's splicing and degradation rates are constant across all cells, we use Equation 38 to evaluate likelihoods, with $\mu_N = \frac{\gamma_g}{\beta_g} \rho_{cg} \ell_c$, $\mu_M = \rho_{cg} \ell_c$, and a shared fit parameter $\alpha_g$. On the other hand, if we relax this assumption, we use the same equation, with $\mu_N = \rho_{cg}^{(N)} \ell_c$ and $\mu_M = \rho_{cg}^{(M)} \ell_c$.

##### S1.2.2 "Cell size" as technical variation

As in Section S1.1.2, we may propose a model of imperfect capture that recapitulates the functional form in *scVI*. Under Bernoulli sampling, the univariate negative binomial model stays negative binomial with the same shape parameter:

$$\begin{aligned}\mu_{cg} &= \frac{\langle K\rangle_{cg}}{\gamma_{cg}} p_c = \frac{\alpha_g \eta_z}{\gamma_z} p_c := \mu^b_{cg} p_c \\ &= \frac{\alpha_g \eta_z}{C\gamma_z} C p_c,\end{aligned} \tag{41}$$

i.e., downsampling amounts to rescaling $\eta_z/\gamma_z$. The *scVI* variables can be interpreted as follows:

$$\begin{aligned}\rho_{cg} &:= \frac{\alpha_g \eta_z}{C\gamma_z}, \\ \ell_c &:= C p_c.\end{aligned} \tag{42}$$

In the two-species case, the same identities hold [37]. The bivariate law is given by Equation 38 with $\mu_N = \mu^b_N p_c$ and $\mu_M = \mu^b_M p_c$. As above, the assumption of a common $p_c$ is not mandatory. Thus, if $\beta$ and $\gamma$ can differ, we find:

$$\begin{aligned}\mu^{(N)}_{cg} &= \alpha_g \frac{\eta_z}{\beta_z} \ell_c = \rho^{(N)}_{cg} \ell_c, \\ \mu^{(M)}_{cg} &= \alpha_g \frac{\eta_z}{\gamma_z} \ell_c = \rho^{(M)}_{cg} \ell_c.\end{aligned} \tag{43}$$

We recapitulate the same functional form under a distinct series of assumptions. To obtain this solution, we used a convenient property by which a Bernoulli-subsampled negative binomial distribution remains negative binomial.

#### S1.3 The negative binomial model and its bursty extensions

Alternatively, the following model can produce a negative binomial stationary distribution of $\mathcal{X}$:

$$\varnothing \xrightarrow{k} B \times \mathcal{X} \xrightarrow{\gamma} \varnothing, \tag{44}$$

i.e., transcription events arrive at rate $k$, and each one induces a *burst* of activity, with the number of produced transcripts $B$ drawn from a geometric distribution on $\mathbb{N}_0$ with mean $b$. In the parlance of Equation 2, the law of $\mathcal{X}$ is negative binomial with shape $\alpha = k/\gamma$ and mean $\mu = kb/\gamma$.

##### S1.3.1 "Cell size" as biological variation

To include dependence on $z$ and the "cell size," while maintaining the assumption of constant shape parameter, we recall that the burst model arises in a particular limit of the two-state telegraph model [15]. The burst frequency $k$ is the rate at which the gene turns on, whereas the burst size $b$ is the average number of molecules produced while the gene is on. The burst size is proportional to the transcription initiation rate, which is in turn controlled by [RNAP]. By choosing the appropriate units and separating out the initiation term to yield $b_{cg} = b_{z,init}[\text{RNAP}]_c$, we obtain

$$
\begin{aligned}
\mu_{cg} &= \frac{k_g b_{cg}}{\gamma_g} = \alpha_g b_{z,init}[\text{RNAP}]_c \\
&= \frac{\alpha_g b_{z,init}}{C} C[\text{RNAP}]_c, \\
\rho_{cg} &:= \frac{\alpha_g b_{z,init}}{C}, \\
\ell_c &:= C[\text{RNAP}]_c.
\end{aligned} \tag{45}
$$

Next, we write down the two-species system:

$$
\varnothing \xrightarrow{k} B \times \mathcal{N} \xrightarrow{\beta} \mathcal{M} \xrightarrow{\gamma} \varnothing, \tag{46}
$$

which induces the a distribution with the following probability-generating function (PGF) [18]:

$$
\begin{aligned}
G(g_N, g_M; k, b, \beta, \gamma) &= \sum_{n,m=0}^{\infty} g_N^n g_M^m P(n, m) \\
&= \exp\left( k \int_0^{\infty} \left[ \frac{1}{1 - bU(u_N, u_M, s; \beta, \gamma)} - 1 \right] ds \right), \\
\text{s.t. } U(u_N, u_M, s; \beta, \gamma) &= \frac{\beta}{\beta - \gamma} u_M e^{-\gamma s} + \left( u_N - \frac{\beta}{\beta - \gamma} u_M \right) e^{-\beta s} \\
\text{and } u_i &:= g_i - 1.
\end{aligned} \tag{47}
$$

As in the generic case, $\mu_N = kb/\beta$ and $\mu_M = kb/\gamma$. The bivariate distribution no longer has a well-defined and unique $\alpha_g$ parameter; for consistency with the "shape of the negative binomial marginal" interpretation, we choose $\alpha := k/\beta$. Equation 47 typically cannot be evaluated in closed form, but may be approximated:

$$
P(n, m; \alpha, \mu_N, \mu_M) = P_{\text{NB}}(n; \alpha, \mu_N) P(m|n; \alpha, \mu_N, \mu_M), \tag{48}
$$

where $P(m|n)$ can be represented by a mixture of negative binomial distributions [19].

We maintain the assumptions of $z$ and "cell size" dependence through $b$, and impose the assumption that $k$, $\beta$, and $\gamma$ are gene-specific and constant. This yields:

$$
\begin{aligned}
\mu_{cg}^{(N)} &= \frac{k_g b_{z,init}}{\beta_g C} = \frac{\alpha_g b_{z,init}}{C} = \rho_{cg} \ell_c, \\
\mu_{cg}^{(M)} &= \frac{k_g b_{z,init}}{\gamma_g C} = \frac{\beta_g}{\gamma_g} \frac{\alpha_g b_{z,init}}{C} = \frac{\beta_g}{\gamma_g} \rho_{cg} \ell_c,
\end{aligned} \tag{49}
$$

where $\beta_g/\gamma_g$ may be fit or estimated from the plug-in method of moments estimate $\overline{M}_g/\overline{N}_g$.

As before, if we suppose that $k$, $\gamma$, and $\beta$ can differ between cell types, we are forced to define two distinct $\rho$ parameters:

$$
\begin{aligned}
\mu_{cg}^{(N)} &= \rho_{cg}^{(N)} \ell_c \text{ s.t. } \rho_{cg}^{(N)} := \frac{k_z b_{z,init}}{C \beta_z}, \\
\mu_{cg}^{(M)} &= \rho_{cg}^{(M)} \ell_c \text{ s.t. } \rho_{cg}^{(M)} := \frac{k_z b_{z,init}}{C \gamma_z}.
\end{aligned} \tag{50}
$$

However, this highly generic form violates admissibility under the *scVI* model: some shape-like parameter should be constant. Maintaining $\alpha_g = k_g/\beta_g$, we yield the following formulation:

$$\begin{aligned} \mu_{cg}^{(N)} &= \rho_{cg}^{(N)} \ell_c \text{ s.t. } \rho_{cg}^{(N)} := \frac{\alpha_g b_{z,init}}{C}, \\ \mu_{cg}^{(M)} &= \rho_{cg}^{(M)} \ell_c \text{ s.t. } \rho_{cg}^{(M)} := \frac{\beta_g}{\gamma_z} \frac{\alpha_g b_{z,init}}{C}. \end{aligned} \tag{51}$$

In sum, if we assume each gene's splicing and degradation rates are constant across all cells, we use Equation 48 to evaluate likelihoods, with $\mu_N = k_g b_{cg}/\beta_g = \frac{\gamma_g}{\beta_g} \rho_{cg} \ell_c$, $\mu_M = \rho_{cg} \ell_c$, and a shared fit parameter $\alpha_g = k_g/\beta_g$. On the other hand, if we relax the assumption of constant $\gamma_g$, we use the same equation, with $\mu_N = \rho_{cg}^{(N)} \ell_c$ and $\mu_M = \rho_{cg}^{(M)} \ell_c$.

##### S1.3.2 "Cell size" as technical variation

Just as in Section S1.2.2, the univariate model stays negative binomial after Bernoulli technical noise:

$$\begin{aligned} \mu_{cg} &= \frac{k_g b_{cg}}{\gamma_g} p_c = \alpha_g b_z p_c, \\ &= \alpha_g \frac{b_z}{C} C p_c, \end{aligned} \tag{52}$$

i.e., the downsampling amounts to rescaling $b$. The *scVI* variables can be interpreted as follows:

$$\begin{aligned} \rho_{cg} &:= \alpha_g \frac{b_z}{C}, \\ \ell_c &:= C p_c. \end{aligned} \tag{53}$$

The two-species model is somewhat more complicated. Specifically, we find that the characteristic takes the following form:

$$\begin{aligned} bU(u_N, u_M, s; \beta, \gamma) &= b p_N u_N e^{-\gamma s} + b p_M u_M \frac{\beta}{\beta - \gamma} \left( e^{-\beta s} - e^{-\gamma s} \right) \\ &= b p_N \left[ u_N e^{-\gamma s} + \frac{p_M}{p_N} u_M \frac{\beta}{\beta - \gamma} \left( e^{-\beta s} - e^{-\gamma s} \right) \right]. \end{aligned} \tag{54}$$

By evaluating $u_N$ and $u_M$ at zero, we can verify that the nascent and mature marginals separately follow the functional form in Equation 47 with burst sizes $bp_N$ and $bp_M$. However, elsewhere, the integrand violates this form: for example, if we fix $\gamma$, there is no way to choose $\beta$ so $p_M/p_N$ will be canceled out at each $s$. This suggests the integral on $s \in [0, \infty)$ – i.e., the stationary PGF – does not match that of the original distribution.

As the current study primarily focuses on the recovery of biophysical, rather than technical parameters, we elide this problem by assuming $p_N = p_M$. This is a coarse approximation that presupposes the sequencing efficiencies of spliced and unspliced RNA are roughly equivalent. It is likely that the relevant parameters can be inferred, or approximated by a simple plug-in estimate [36].

Thus, under the assumption of $p_N = p_M = p$, we yield the law given in Equation 47 with the $b$ variable set to $bp_c$. Enforcing a constant $\alpha_g = k_g/\beta_g$, we yield the following results:

$$\begin{aligned} \mu_{cg}^{(N)} &= \alpha_g \frac{b_z}{C} C p_c = \rho_{cg}^{(N)} \ell_c, \\ \mu_{cg}^{(M)} &= \alpha_g \frac{\beta_g}{\gamma_z} \frac{b_z}{C} C p_c = \rho_{cg}^{(M)} \ell_c, \end{aligned} \tag{55}$$

which recapitulates the results in the previous section.

#### S1.4 The zero-inflated negative binomial model and its multi-state interpretation

The default *scVI* generative model is zero-inflated negative binomial (ZINB), which has the following functional form:

$$\begin{aligned} P(x; \alpha, \mu, \pi) &= \pi P_{\text{NB}}(x; \alpha, \mu) \text{ when } x > 0, \text{ and} \\ &= (1 - \pi) + \pi P_{\text{NB}}(0; \alpha, \mu) \text{ when } x = 0. \end{aligned} \tag{56}$$

The quantity $\pi \in (0, 1)$ is the zero-inflation probability, which is computed using a neural network in the standard *scVI* pipeline.

Despite its popularity, the zero-inflated model is somewhat contentious [39], with a fair amount of evidence suggesting that zero-inflation is not necessary to summarize the distributions of molecule counts in single-cell RNA sequencing data [40]. Our previous mechanistic studies in relatively homogeneous cell types [36, 37] have not implicated zero-inflation as a significant effect.

More problematically, zero-inflation appears to be a purely *ad hoc* mathematical tool, and does not describe any specific biophysical process. We are aware of one report that attempts to motivate it [41]. However, we find this attempt to be unrealistic, as it conflates biological processes with experimental ones: the report describes a *biological* state that leads to *technical* non-identifiability, without motivating a chemical mechanism.

In principle, we *can* write down a system that yields a zero-inflated negative binomial stationary distribution, a simple three-state model [42]:

$$\begin{aligned} &\mathcal{G}_0 \underset{k_{10}}{\overset{k_{01}}{\rightleftarrows}} \mathcal{G}_1, \\ &\mathcal{G}_1 \underset{k_{21}}{\overset{k_{12}}{\rightleftarrows}} \mathcal{G}_2, \\ &\mathcal{G}_2 \xrightarrow{k_{init}} \mathcal{G}_2 + \mathcal{X}. \end{aligned} \tag{57}$$

We omit the downstream processes, which are generic and immaterial. Ostensibly, if $k_{01} = 0$, this reduces to the two-state model. If $k_{21}, k_{init} \to \infty$ such that $k_{init}/k_{21}$ is finite, this further reduces to the bursty model with burst size $b = k_{init}/k_{21}$ and burst frequency $k = k_{12}$. In this limit, the instantaneous probability of being in state $\mathcal{G}_2$ vanishes.

If we impose $k_{10}, k_{01} \ll k_{12}$, but $k_{10}/k_{01} \sim \mathcal{O}(1)$, this system reduces to a two-component mixture, with $\pi = \frac{k_{10}}{k_{01}+k_{10}}$ giving the probability that the gene is in state $\mathcal{G}_1$. This recapitulates the functional form of the ZINB model. In sum, this probability law can emerge from a purely biological description of a system with two long-lived steady states; one of these steady states is inactive, whereas the other exhibits bursts of activity. Analogous models have previously been proposed for co-regulation of gene loci [42], and can be leveraged to represent patterns of co-bursting [43]. For simplicity, we do not discuss this extension further, but the associated derivations would be largely analogous to those in Section S1.3.

#### S2 *biVI* constitutive generative process

*biVI*'s generative process for the constitutive hypothesis models expression values of $x_{cn}$ and $x_{cm}$ of nascent and mature counts, respectively, in cell $c$ as:

$$\begin{aligned}
z_c &\sim \text{Normal}(0, I) \\
\ell_c &\sim \log\text{normal}(\ell_\mu, \ell_{\sigma^2}) \\
\rho_{cg}^{(N)}, \rho_{cg}^{(M)} &= f(z_c, s_c) \\
\mu_{cn}, \mu_{cm} &= \rho_{cg}^{(N)}\ell_c, \rho_{cg}^{(M)}\ell_c \\
x_{cn}, x_{cm} &\sim P_{\text{const}}(n, m; \mu_{cn}, \mu_{cm}) = \text{Poisson}(n; \mu_{cn})\text{Poisson}(m; \mu_{cm}),
\end{aligned} \tag{58}$$

with modeling choices as described in Section 2.4, excepting the need of the jointly optimized $\alpha$. Relative splicing rate $\beta/k$ and relative degradation rate $\gamma/k$ can be recovered according to the following conversions:

$$\begin{aligned}
\beta/k &= \frac{1}{\mu_{cm}} \\
\gamma/k &= \frac{1}{\mu_{cn}},
\end{aligned} \tag{59}$$

where we again set $k = 1$.

#### S3 Extrinsic generative process

*biVI*'s generative process for the extrinsic hypothesis models expression values of $x_{cn}$ and $x_{cm}$ of nascent and mature counts, respectively, in cell $c$ as:

$$\begin{aligned}
z_c &\sim \text{Normal}(0, I) \\
\ell_c &\sim \log\text{normal}(\ell_\mu, \ell_{\sigma^2}) \\
\rho_{cg}^{(N)}, \rho_{cg}^{(M)} &= f(z_c, s_c) \\
\mu_{cn}, \mu_{cm} &= \rho_{cg}^{(N)}\ell_c, \rho_{cg}^{(M)}\ell_c \\
x_{cn}, x_{cm} &\sim P_{\text{extrinsic}}(n, m; \mu_{cn}, \mu_{cm}),
\end{aligned} \tag{60}$$

with modeling choices as described in Section 2.4 and $P_{\text{extrinsic}}(n, m; \mu_{cn}, \mu_{cm})$ defined as Equation 11. Here, we set $\alpha = \langle K \rangle$, or the mean of the transcription rate distribution of Equation 11. Relative splicing rate $\beta/k$ and relative degradation rate $\gamma/k$ can then be recovered according to the following conversions:

$$\begin{aligned}\beta/k &= \frac{\alpha}{\mu_{cn}} \\ \gamma/k &= \frac{\alpha}{\mu_{cm}},\end{aligned} \tag{61}$$

where we again set $k = 1$.

### S4 Network architecture

The *biVI* architecture used for all figures and analyses in this study employs two encoders of 3 fully connected, 128 node layers to map from input space $\mathbb{R}^{2N_G}$, where $N_G$ is the number of genes, to latent means $z_\mu \in \mathbb{R}^{10}$ and latent standard deviations $z_\sigma \in \mathbb{R}^{10}$. The decoder, comprising 3 fully connected layers of 128 nodes, maps from sampled latent space $z_n \in \mathbb{R}^{10}$ back to the original input space $\mathbb{R}^{2N_G}$. There are ReLU activation functions between each layer of the encoder networks and decoder network. Furthermore, a softmax is applied to the output layer of the decoder to produce normalized expression levels $\rho_{cg}^{(N)}$ and $\rho_{cg}^{(M)}$ for nascent and mature expression levels, respectively. $\rho_{cg}^{(N)}$ and $\rho_{cg}^{(M)}$ are multiplied by sampled sequencing depth $\ell_c$ before conditional likelihoods (reconstruction errors) are calculated. For the bursty and extrinsic models, the architecture includes parameters $\alpha \in \mathbb{R}^{N_G}$ that are updated during inference. For the constitutive model, only nascent and mature means parameterize the conditional likelihood, so $\alpha$ is unused.

### S5 Connecting the models to transcriptome data

Thus far, we have used "nascent" and "mature" as shorthands for the discrete species in a two-stage model of RNA processing. In other words, Equation 5 is axiomatic for this nomenclature. We have gone one step further and named the rate of conversion $\beta$ the *splicing rate*, explicitly identifying the nascent species with unspliced RNA and the mature species with spliced RNA. This identification is a modeling decision that elides the considerable simplification of biological complexities. In the current section, we elaborate on the assumptions.

In the field of microbiology, "nascent" RNA is often, but not always, used to characterize the mRNA molecules in the process of synthesis, associated to a DNA strand via an RNA polymerase complex [44–47]. In this framework, the "mature" transcriptome is simply the complement of the nascent transcriptome, i.e., all molecules that are not chemically associated to a DNA strand. Therefore, the canonical definition of "nascent" RNA is equivalent to *transcribing* RNA, which is a polymeric structure with a particular sequence.

Transcribing RNA can be observed directly through electron micrography [46]. However, more typically, it is investigated through more or less direct experimental proxies that can be scaled to many genes and cells at a time. In the single-cell fluorescence subfield, DNA or membrane staining can be used to identify bright spots localized to the nucleus, which is treated as signal from RNA at the transcription site [25, 48]; this signal may include contributions from RNA incidentally, or mechanistically, retained at a DNA locus [45]. In this strategy, "nascent" molecules are DNA-associated. Alternatively, and more commonly, transcribing molecules have been studied by using probes targeted to the 5′ and 3′ regions [42, 49–51], or to intronic and exonic regions [52–55]. In this strategy, "nascent" molecules contain a particular region, either synthesized earlier or removed later in the RNA life-cycle.

The use of intron data as a proxy for active transcription is reminiscent of, but distinct from sequence census [56] strategies that directly study RNA sequences. These strategies, in turn, typically use chemical methods to enrich for newly transcribed RNA. For example, Reimer et al. isolate chromatin, then deplete sequences that have been post-transcriptionally poly(A) tailed [57]. Analogously, Drexler et al. use 4-thiouridine (4sU) labeling to enrich for newly synthesized molecules [58]. These approaches may produce conflicting results; for example, introns may be rich both in poly(A) handles [1] and 4sU targets [57], giving rise to obscure technical effects. Therefore, these "processed" or "temporally labeled" proxies are coarsely representative of transcriptional dynamics, and their quantitative interpretability is unclear as of yet.

The sequence content may be used more directly, by conceding that DNA association or localization are not easily accessible by sequence census methods, and treating splicing *per se*. This approach has a fairly long history. Intronic quantification has been used to characterize transcriptional mechanisms in microarray datasets [59], and to characterize differentiation programs in RNA sequencing [60, 61]. In single-cell RNA sequencing, intronic content has been leveraged to identify transient behaviors from snapshot data [1], albeit with some outstanding theoretical concerns and caveats [62]. Briefly, it is, in principle, possible to coarsely classify molecules with intronic content as "unspliced" or "pre-mRNA" and aggregate all others as "spliced," "mature," or simply "mRNA." By applying this binary classification, and defining a simple first-order model of splicing, data may be successfully fit. The quantification of transcripts so classified is a relatively straightforward genomic alignment problem. The multiple available implementations [1, 2, 63, 64] tend to disagree on the appropriate assignment of ambiguous sequencing reads [62, 64], obscuring a more fundamental problem: the binary classification is mechanistically limited [58, 65–67], and it is likely that detailed splicing graph models will be necessary in the future [43, 62]. The focus on sequence is yet another step removed from the transcriptional dynamics, particularly since some of the splicing processes occur after transcriptional elongation is complete [68]. However, in spite of its limitations and assumptions, the simple Markovian two-stage model has been successful in the past [36, 37]. Mechanistic evidence does not suggest that, e.g., deterministic delayed elongation is necessary to represent "unspliced" distributions [69]: under this model, we appear to be able to treat splicing and degradation as Markovian, without representing the elongation process at all, and obtain reasonable fits to the data. Somewhat surprisingly [70], the geometric-Poisson distribution, which describes bursty transcription coupled to deterministic elongation [71], is a particularly poor fit for unspliced RNA counts [69].

Adding yet more complexity to the modeling, "mature" – whether "off-template," "spliced," or "processed" – molecules are not immediately available for degradation; first, the process of nuclear export must take place. Studies that presuppose access to imaging data tend to model it explicitly [18, 25, 72–74]. However, this approach has not been applied in sequencing assays, as current technologies do not distinguish nuclear and cytoplasmic molecules. Furthermore, comparisons of paired single-cell and single-nucleus datasets are hampered by the limited characterization of the noise sources in the latter technology. Yet again, we have generally found that omitting this effect in single-cell data produces acceptable fits [36, 37, 69].

Pending the development of more sophisticated sequencing and alignment technologies, as well as the implementation of tractable models of biology, the data exploration portion of our study focuses on the "spliced" and "unspliced" matrices generated by *kallisto* |*bustools* [2]. This choice is a compromise, and we adopt it after considering the following factors:

- Availability of quantification workflows: spliced and unspliced matrices are straightforward to generate.
- Model tractability: the two-stage models can be evaluated [19]; more sophisticated models require new algorithms to be integrated with variational autoencoders.
- The scope of sequencing data: single-cell protocols do not yet give access to sub-cellular information, so inference of elongation or nuclear retention dynamics is acutely underspecified.
- Self-consistency and compatibility: *a priori*, we seek to recapitulate the data types (spliced RNA counts) and distributions (negative binomial and Poisson) already implemented in *scVI*, leaving improvements to further investigation.

We use the terms "nascent" and "mature" to identify the unspliced and spliced RNA matrices. This choice of nomenclature is deliberate. Although it somewhat conflicts with the established microbiology literature, this terminology is intended to emphasize the models' generality, both here and elsewhere [9, 19, 75]. The two-stage Markovian process is axiomatic. The specific identities assigned to the mathematical objects may range beyond counts identified by sequence census methods. They may represent the discretized and subtracted intensities of 3', 5', intron, or exon fluorescent probes, the counts of molecules within and outside the nuclear envelope, or polymerase counts obtained by micrography. Therefore, the terminology should be taken in the sense used for similar non-delayed models in [70, 72, 76, 77].

### S6 Simulations

#### S6.1 Simulated data

To validate our implementation of *biVI*, and to understand potential pitfalls inherited from the standard *scVI* autoencoder workflow, we generated ground truth data by simulation. Simulated ground truth is limited in important ways, as it omits known and unknown sources of biological and technical variability. However, to begin to understand the limitations of integrating descriptive autoencoders with mechanistic models, it is helpful to have simple and well-understood simulated datasets. By judiciously choosing simulation scenarios, we can characterize the relative and absolute accuracy of the procedures under ideal-case conditions, which describe a natural upper bound on their performance.

For each of the three models (constitutive, extrinsic, and bursty), we simulated a data set of nascent and mature RNA counts for 2,000 genes across 10,000 cells, spread across 20 cell types. The cell types were distinguished by low-magnitude variation in all genes' parameters, as well as higher-magnitude upregulation of expression in a small set of cell type-specific marker genes. In the extrinsic noise model, upregulation was effected by increasing the transcription rate scale $\eta$. In the analogous constitutive case, it was effected by increasing the transcription rate $k$. In the bursty model, it was effected by increasing the burst size $b$.

#### S6.2 Generating parameters

For all models, parameters were generated for 2,000 genes across 20 cell types. First, baseline $\log_{10}$ model parameters were drawn from multivariate normal distributions for all 2,000 genes.

For the bursty model, mean burst size $b$, relative splicing rate $\beta/k$, and relative degradation rate $\gamma/k$ were sampled from to the following trivariate normal distribution:

$$\begin{pmatrix} \log_{10} b \\ \log_{10} \frac{\beta}{k} \\ \log_{10} \frac{\gamma}{k} \end{pmatrix} \sim N \left[ \begin{pmatrix} 0.2 \\ 0.5 \\ 0.4 \end{pmatrix}, \begin{pmatrix} 0.36 & 0.144 & 0.24 \\ 0.144 & 0.09 & 0.12 \\ 0.24 & 0.12 & 0.5 \end{pmatrix} \right].$$

For the constitutive model, parameters for relative splicing rates $\beta/k$ and relative degradation rates $\gamma/k$ were generated from the following bivariate normal distribution:

$$\begin{pmatrix} \log_{10} \frac{\beta}{k} \\ \log_{10} \frac{\gamma}{k} \end{pmatrix} \sim N \left[ \begin{pmatrix} 0.497 \\ 0.711 \end{pmatrix}, \begin{pmatrix} 0.318 & 0.062 \\ 0.062 & 0.237 \end{pmatrix} \right].$$

Finally, for the extrinsic model, the negative binomial shape parameter $\alpha$, relative splicing rate $\beta/\eta$, and relative degradation rate $\gamma/\eta$ were generated from the following trivariate normal distribution:

$$\begin{pmatrix} \log_{10} \alpha \\ \log_{10} \frac{\beta}{\eta} \\ \log_{10} \frac{\gamma}{\eta} \end{pmatrix} \sim N \left[ \begin{pmatrix} 2.53 \\ 0.31 \\ 0.25 \end{pmatrix}, \begin{pmatrix} 1.01 & 0.26 & -0.08 \\ 0.26 & 0.49 & 0.11 \\ -0.08 & 0.11 & 0.21 \end{pmatrix} \right].$$

The parameter distribution for the bursty and extrinsic models were derived from *Monod* fits to a cell type dataset (assuming no technical noise) [37]. The parameter distribution for the constitutive model was derived from a joint distribution of nascent and mature means.

After creating the baseline, we added random noise, distributed per $\epsilon \sim N(0, 0.05)$ to all $\log_{10} b$ and $\log_{10} \frac{\gamma}{k}$ for the bursty model, and $\log_{10} \frac{\beta}{k}$ and $\log_{10} \frac{\gamma}{k}$ for the constitutive and extrinsic models per cell type. Therefore, each cell type had slightly different parameters for non-marker genes. For consistency with default *scVI* learning assumptions, we did not change the relative splicing rate $\frac{\beta}{k}$ of the bursty model and the inverse dispersion parameter $\alpha$ of the extrinsic model: though different across genes, they were identical across all cell types.

Next, we imposed more realistic cell type structure by selecting marker genes. The number of marker genes per cell type was chosen for each cell type by drawing from a Poisson distribution with a mean of 45. Marker genes were non-overlapping. In the bursty model, marker gene burst sizes were augmented by $\Delta \sim N(1.0, 0.1)$ in the relevant cell type. In the constitutive model, marker gene parameters were modified by increasing the rate of transcription $k$, effected by subtracting $\Delta \sim N(1.0, 0.1)$ from each $\log_{10} \beta/k$ and $\log_{10} \gamma/k$. In the extrinsic model, marker gene parameters were modified by increasing the scale of transcription $\eta$, effected by subtracting $\Delta \sim N(1.0, 0.1)$ from each $\log_{10} \beta/\eta$ and $\log_{10} \gamma/\eta$.

#### S6.3 Sampling count data

10,000 cells spread across the 20 cell types were simulated for each model. The number of cells per cell type was assigned using a Dirichlet-multinomial distribution: a probability vector $\mathbf{p}$ was drawn from a Dirichlet distribution parameterized by a vector of 20 equal elements, then cell types for each of the 10,000 cells were assigned using a multinomial distribution with probabilities $\mathbf{p}$. Nascent and mature counts for each of the 10,000 cells were then sampled from model distributions (Equation 13 for bursty data, Equation 7 for constitutive data, and Equation 11 for extrinsic data) using parameters specific to the cell's assigned cell type.

##### S6.4 Fitting simulated data

We fit the three simulated data sets using *biVI* with the three generative physical models, as well as standard *scVI* with the negative binomial distribution as its conditional likelihood. All models were trained on 7,200 cells with 800 validation cells for 400 epochs with a learning rate of 0.001. The encoder and decoder networks consisted of 3 layers with 128 nodes each, with a latent space of dimension 10. As in standard *scVI*, a standard normal prior was used for the latent space.

##### S6.5 Performance metrics on simulated data

Performance of the trained models was tested on 2,000 held-out cells for each simulated data set. In order to characterize the absolute reconstruction of ground truth cell types, the mean squared error (MSE) between *biVI* inferred means, $\mu_N$ and $\mu_M$, and ground truth simulated means was calculated for each cell. Furthermore, we assessed how well different models reduced cells of the same cell type to similar latent spaces while maintaining good separation of cells in different cell types. To compare clustering accuracy for different models, we applied several metrics to the latent spaces obtained for testing data, using simulated ground truth cell type as cluster assignment. We calculated average intra-cluster distance (ICD), or the Euclidean distance between each cell's latent representation and the mean of its assigned cluster in the latent space averaged over all cells. We also calculated average intra-cluster variance (ICV), or the variance of ICDs for each cell in a cluster averaged over all clusters. Inter-cluster distance was also calculated: distance between cluster means averaged over all pair-wise clusters (excluding the cluster's distance to its own mean). Finally, scaled inter-cluster distance, or inter-cluster distance divided by average intra-cluster distance, was found. We also report nearest neighbor percentages (Figures S3, S4, S5), or the percent of the $n_\kappa$ nearest neighbors in the same cell type as each cell, where $n_\kappa$ is the number of cells in the given cell's cell type. A quantitative description of clustering metrics is provided in Section S7. Broadly speaking, these metrics characterize the methods' utility for discovering discrete cell populations using typical clustering algorithms, which perform best when low-dimensional clusters are relatively condensed, homogeneous, and reflect the high-dimensional structure.

##### S6.6 Kullback-Leibler divergence between simulated and reconstructed distributions

We compare *biVI* and *scVI*'s ability to accurately reconstruct ground truth gene distributions (posterior predictive distributions) by calculating the truncated Kullback-Leibler divergence (KLD) between ground truth simulated distributions and reconstructed (or, posterior predictive) distributions for all genes in all cell types. We reconstruct gene and cell type specific *biVI* and *scVI* distributions $\hat{P}(n, m)$ as described above in Section 3. We also calculate true distributions $Q_{\kappa g}(n, m; \theta_{\kappa g})$ using the gene and cell type specific ground truth simulated parameters $\theta_{\kappa g}$ (parameter generation and sampling are described in Section S6). We calculate both true and reconstructed model (*biVI* or *scVI*) distributions over a 50 by 50 grid of nascent and mature values, then normalize both to sum to 1.0 over the grid by dividing all probabilities by their sum over the grid. Normalization ensures that KL divergences are well-defined. We then calculate the KL divergence over the restricted grid:

$$\text{KLD} = \sum_{m=0}^{49} \sum_{n=0}^{49} \hat{P}_{\kappa g}(n, m) \times \log \frac{\hat{P}_{\kappa g}(n, m)}{Q_{\kappa g}(n, m; \theta_{\kappa g})}. \tag{62}$$

We note that conditional data likelihood distributions and true, marginal data distributions are *not* necessarily of the same form (for example, if the conditional data likelihood distribution is negative binomial for a gene in a cell type, the marginal distribution of the gene in that cell type is not necessarily negative binomial). The above KLD calculated for simulated data, then, in which we reconstruct marginal data distributions by treating the conditional data likelihood distribution as having the same form as the true marginal distribution from which simulated data was sampled, is not a perfect comparison but still allows a relative of estimate of how well different models can reconstruct known ground truth distributions.

##### S6.7 Simulated results

For all simulated datasets, the negative log-likelihood of data, or reconstruction loss, of the 2,000 held-out testing cells was lower for *biVI* models that used the simulated data's generative distribution as the conditional likelihood than for *scVI* models that used independent negative binomial distributions as conditional likelihoods (Figures S3j, S4j, and S5j). For example, training *biVI* with the bursty generative model on a bursty simulated data set resulted in a reconstruction loss of 3,885.8 while that of *scVI* trained with negative binomial data distributions on the same data 3,953.6. Furthermore, the Kullback-Liebler divergence (KLD) between true and reconstructed count distributions (Section 3), separately computed for each cell type, was lower for *biVI* models with the simulated generative distribution than for *scVI* (Figures S3h, S4h, and S5h). On bursty simulated data, the average KLD between cell type's true and reconstructed distribution was 0.014 for *biVI* with the bursty generative model and 0.0212 for *scVI* (Figure S3h). These results suggests that *biVI* can better capture true underlying data distributions than *scVI*, providing a potential route to mechanistic model selection for observed data based on recapitulation of empirical data distributions.

A primary use of *scVI* is to reduce the dimensionality of similar cells, such as multiple observations of the same cell type, to similar low-dimensional latent representations, removing noise and redundancies. To characterize how well cell types are preserved in the latent space, we applied several clustering metrics (Section S7) to the latent representation of 2,000 cells for each model encoded by *biVI* models or *scVI*, using the ground truth cell types as cluster assignments (Figure S3a-e,g, S4a-e,g, and S5a-e,g). For bursty data, the average percent of latent-space nearest neighbors in the same cell type as each cell, computed over all cells, was similar for *biVI* with the bursty generative model (90.0%) and *scVI* (90.4%) (Figure S3g). The silhouette score, a measure of how similar each cell is to other cells in its cluster compared to other clusters, achieved analogous performance, yielding 0.383 for *biVI* and 0.387 for *scVI* (Figure S3e). Finally, scaled inter-cluster distance (average inter-cluster distance divided by average intra-cluster distance, with larger values corresponding to better cluster separation), was 2.75 for *biVI* and 2.60 for *scVI* (Figure S3d). These clustering metrics are evidence that cells can be similarly well represented in a low dimensional space using *biVI* and *scVI*, and that *biVI* modifications do not impede the process of variational inference or downstream analyses.

##### S6.8 Simulated marker gene detection

To determine how well *biVI* and *scVI* models discover ground-truth marker genes, we performed Bayesian hypothesis testing over a range of $log_2$ fold change and Bayes factor thresholds (see Section 2.8) on the bursty simulated dataset. cell types had between 37 and 59 marker genes out of 2,000 total genes; for each cell type, we identified differential genes between that cell type and all others. To facilitate comparison, we performed testing on unspliced and spliced mean values, storing lists of genes identified as differential in unspliced mean, spliced mean, and "either," or the union of the unspliced and spliced differential genes.

We varied the $log_2$ fold change (LFC) effect size (denoted $T$ in Section 2.8) between 0.2 and 2.0 with a constant Bayes factor threshold of 0.7 (as is default in *scVI*). We also varied log Bayes factor threshold between 0.2 and 2.0, holding LFC effect size $T$ at a constant value of 0.5.

For each Bayes factor calculation, we drew 10 samples from the posterior of each cell in the compared group of cells and permuted parameters for a total of 10,000 comparisons.

We calculated specificity, or true positive rate, defined as the number of true positives (marker genes marked as differentially expressed) divided by the total number of marker genes, and specificity, or true negative rate, defined as the number of true negatives (non-marker genes not marked as differentially expressed) divided by the total number of non-marker genes.

$$\text{Sensitivity} = \frac{\text{TP}}{\text{TP} + \text{FN}} \tag{63}$$

$$\text{Specificity} = \frac{\text{TN}}{\text{TN} + \text{FP}} \tag{64}$$

for unspliced, and spliced means. We took the average and standard deviation of sensitivity and specificity values over the 20 cell types to display in Figure S7.

Figure S7a shows that the bursty *biVI* model discovers almost all marker genes, with a sensitivity near perfect, even as LFC threshold is increased, while other methods fail to identify true marker genes (sensitivity falls to 0.0). However, this does not lead to a dramatic decrease in sensitivity for *biVI*: while other models are more specific, or correctly exclude non-marker genes over increasing LFC values, *biVI* also excludes most non-marker genes (Section S7b). The same trends hold for increasing the Bayes factor threshold while holding the LFC threshold constant: *biVI* correctly identifies marker genes with a high specificity and sensitivity, while other methods fail to identify marker genes (Figure S7c,d).

#### S7 Benchmarking metrics

The following clustering metrics were calculated for simulated data and Allen experimental data on the latent spaces produced by each model. Ground truth simulated cell type was used as cluster assignment for simulated data, and well-annotated cell subclass assignment for Allen datasets. Let $N_C$ be the total number of cells and $J$ be the number of clusters. Let $z_c \in \mathbb{R}^d$ be the low-dimensional representation of cell $c$, $n_\kappa$ be the number of cells in each cluster $\kappa$ (such that $\sum_\kappa n_\kappa = N_C$), and $\mu_\kappa \in \mathbb{R}^d$ be the mean of low-dimensional representations of cells in cluster $\kappa$.

**Average intra-cluster distance (average ICD)** Average intra-cluster distance was found by calculating the Euclidean distance between each cell's low-dimensional representation and its cluster mean's low-dimensional representation. First, the average intra-cluster distance for each cluster $\kappa$ was found:

$$\text{ICD}_{c_\kappa} = \|z_{c_\kappa} - \mu_\kappa\|_2, \tag{65}$$

$$\overline{\text{ICD}_\kappa} = \frac{1}{n_\kappa} \sum_{c_\kappa=1}^{n_\kappa} \text{ICD}_{c_\kappa} \quad \forall\ c_\kappa \in \kappa. \tag{66}$$

Then, the average over all clusters was computed:

$$\text{average ICD} = \frac{1}{J} \sum_{\kappa=1}^{J} \overline{\text{ICD}_\kappa}. \tag{67}$$

For clustering on the latent space, ideally the underlying cell types should be fairly "condensed", with a low average ICD.

**Average intra-cluster variance (average ICV):** The intra-cluster variance was calculated for each cluster $\kappa$ as the variance in intra-cluster distances (ICDs) for all cells in cluster $\kappa$:

$$\text{ICV}_\kappa = \frac{1}{n_\kappa} \sum_{c_\kappa=1}^{n_\kappa} (\text{ICD}_{c_\kappa} - \overline{\text{ICD}_\kappa})^2 \quad \forall\ c_\kappa \in \kappa. \tag{68}$$

Then, the average over all clusters was computed:

$$\text{average ICV} = \frac{1}{J} \sum_{\kappa=1}^{J} \text{ICV}_\kappa. \tag{69}$$

For clustering on the latent space, ideally the average ICV should be low: having a high variation in spread tends to violate the assumptions of typical algorithms.

**Average inter-cluster distance (average INCD):** The distance between each cluster mean and all other cluster means was found. The average was computed over all these distances:

$$\text{average INCD} = \frac{1}{J^2 - J} \sum_{y=1}^{J} \sum_{\substack{x=1 \\ x \neq y}}^{J} \|\mu_x - \mu_y\|_2. \tag{70}$$

For clustering on the latent space, ideally the average INCD should be relatively high, to facilitate discrimination between populations.

**Scaled inter-cluster distance (INCD):** The scaled inter-cluster distance was found by dividing the average inter-cluster distance by the average intra-cluster distance:

$$\text{scaled INCD} = \frac{\text{average INCD}}{\text{average ICD}}. \tag{71}$$

As with the average INCD, ideally this quantity to be relatively high.

**Silhouette score:** The silhouette score was calculated as follows:

$$\frac{b-a}{\max(a,b)},$$

where $a$ is the average intra-cluster distance and $b$ is the average nearest-cluster distance, or mean of the distance between each sample and the nearest cluster that it is not a part of. Silhouette score was implemented using the function `sklearn.metrics.silhouette_score` [78]. Ideally the silhouette score should be high, to accentuate the differences between clusters.

**Nearest neighbor percentage:** Nearest neighbor percentages were calculated for each cell by dividing the number of the $n_\kappa$ nearest cells (in low-dimensional space) in the same cluster $\kappa$ as the cell by the total number of cells, $n_\kappa$, in that cluster.

The computation was implemented using `sklearn.neighbors.KNeighborsClassifier` [78]. Ideally the nearest neighbor percentage should be high, as it helps interpret neighborhood relationships in the latent space as cell type similarity.

**Mean squared error:** The mean squared error (MSE) between reconstructed parameters and ground truth parameters over all genes was calculated. Ideally this quantity should be low, as it confirms that the autoencoder is not merely learning distribution shapes (potentially overfitting or underfitting due to distributional degeneracies), but obtaining quantitative information about the parameter values.

#### S8 Conditional parameter interpretation

In the Bayesian framework, the inferred parameters are conditional on a single sampling from a cell's approximate posterior distribution for its latent representation $z$. The parameters of the marginal data distribution depend on integrating over the posterior distribution.

We explored how variation in the approximate posterior introduces variation in inferred conditional parameters by generating 50 Monte Carlo samples from 30 randomly chosen cell's posteriors, then decoding to produce parameters for a randomly chosen gene for each cell. Figure S8 shows results for sampling from *biVI* with the bursty generative model and *scVI* with independent negative binomial conditional likelihoods trained on Allen data as described in Section 2.7). Normalized parameters, or those not yet multiplied by sequencing depth, are shown in Figure S8a for *biVI* and Figure S8b for *scVI* with their post-scaling counterparts in Figure S8c for *biVI* and Figure S8d for *scVI*. Overlap of conditional parameters between gene-cell pairs for *scVI* nascent and mature mean parameters is apparent, with better separation of *biVI* parameters. We also report the average of decoded parameters over the 50 samples from the posterior in Figure S8e-g: average values tend to converge after few ($< 10$) samples, with initial relationships between conditional parameters preserved (i.e., if one conditional parameter is larger than another at the first sampling, its average value is still larger after many samples). This suggests that the *relative* values of inferred conditional parameters can be reasonably compared and interpreted as reflecting some true differences in underlying marginal data distributions.

#### S9 Linear decoder training and results

A single linear layer can be used as a decoder in place of a multi-layer network with non-linear activation functions to directly link latent variables with inferred gene parameters. To demonstrate this, we trained *biVI* with the bursty generative model on the processed Allen dataset, sample B08, with a linear decoder. We used 4,622/513/1,283 cells for training/validation/testing. The model consisted of an encoder with 3 hidden layers of 128 hidden units with ReLU activation functions, a latent space of 10 dimensions, and a linear decoder layer of 128 nodes with no bias added. We trained for 400 epochs with a learning rate of $10^{-5}$. We found that higher learning rates resulted in undefined gradient calculations: decreasing learning rates led to better stability during training.

Figure S9a-d show that effects of latent dimensions on genes can be found by examining the weights of the linear decoder. A higher weight between a latent dimension and a gene indicates that the latent dimension is positively correlated with the nascent or mature mean expression of that gene, and a lower weight indicates negative correlation. Further, genes can be grouped according to which latent factor most greatly increases their inferred nascent or mature mean value, revealing potentially co-regulated "gene modules" [27]. For now, nascent and mature values are treated separately, as the linear decoder multiplies latent variables by separate weights to produce nascent and mature means (Figure S9a-c). For example, we grouped genes into 10 modules based on the weights in the linear decoder for nascent means. Genes whose nascent means were most greatly increased by latent dimension $z_i$ was placed in group $i$ for all $i \in [0, 9]$ latent dimensions. Then, we performed a Gene Ontology (GO) enrichment analysis for each group of genes to discover enriched biological processes [79]. We performed the analysis with species *Mus musclus* using all genes in the database as a reference list [79]. We plotted the fold enrichment and significance value for the top biological process discovered for each group (Figure S9d) to show that latent dimensions can be used to separate sets of co-regulated gene modules that function in relevant physiological pathways. While we show results for nascent means, similar analyses could be done for weights learned for mature means, or the overlap of nascent-affecting and mature-affecting grouped genes.

We also demonstrate in Figure S9f and g that latent representations learned using a linear decoder can capture capture cell type structure. Nearest neighbor fraction and scaled inter-cluster distance (calculated according to Section S7) were calculated for all cells in the full expression space (using nascent and mature means), in the 10-dimensional latent space obtained using a *biVI* bursty model with a 3-layer non-linear decoder, and in the 10-dimensional latent space obtained using a *biVI* bursty model with a linear decoder. While latent representations obtained using a non-linear decoder have higher nearest neighbor percentages and average scaled inter-cluster distance (higher values indicating better clustering) than for latent representations obtained using a linear decoder, the linear decoder latent representations and full expression matrix cluster cell subclasses comparably well.

#### S10 Supplementary Figures

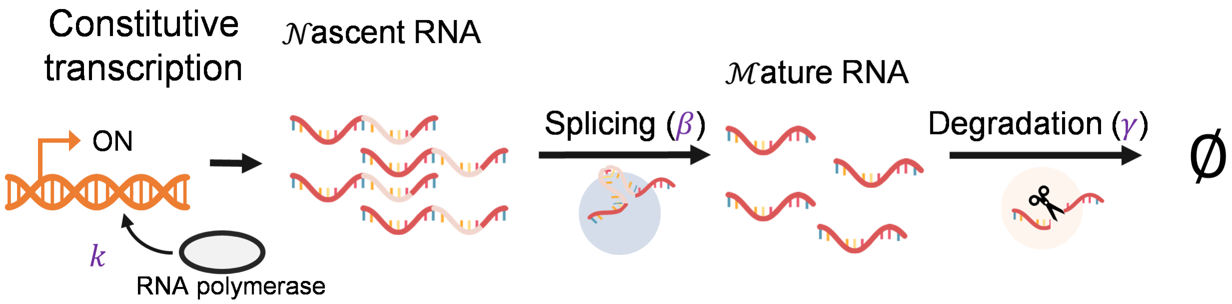


Figure S1: Diagram of constitutive transcription. Transcription occurs at constant rate $k$ to produce nascent RNA molecules. Next, splicing occurs according to a Poisson process with rate $\beta$ to produce mature RNA molecules, which are degraded according to a Poisson process with rate $\gamma$.

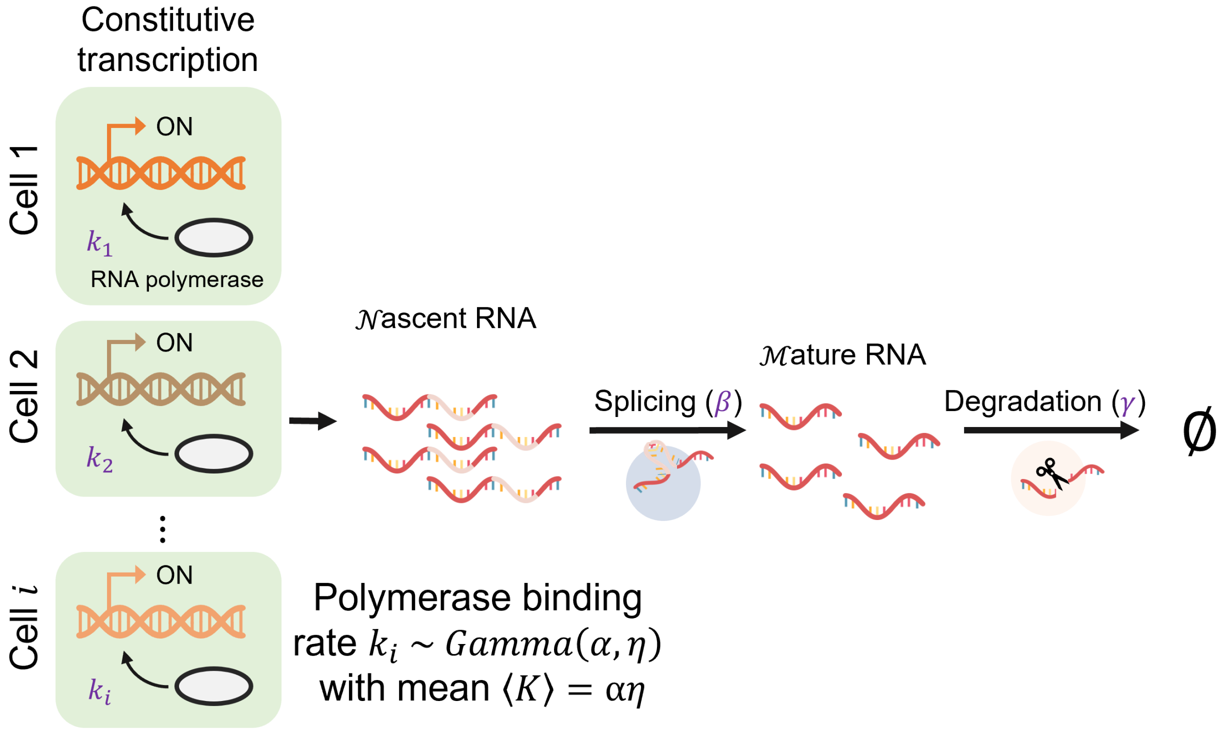


Figure S2: Diagram of the extrinsic model of transcription. Transcription across cells of the same cell type occurs according to a Poisson process with cell-specific rates $k \sim Gamma(\alpha, \eta)$ to produce nascent RNA molecules. Next, splicing occurs according to a Poisson process with rate $\beta$ to produce mature RNA molecules, which are degraded according to a Poisson process with rate $\gamma$.

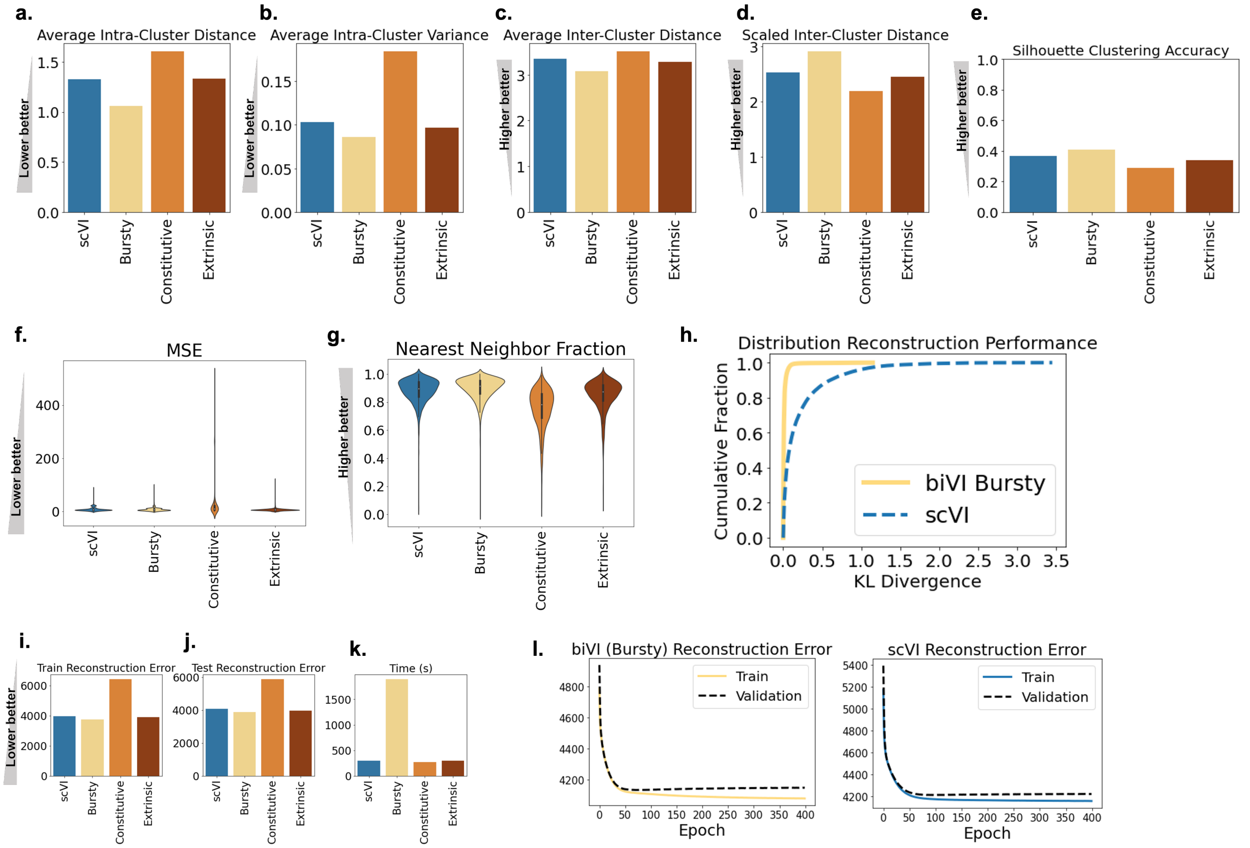


Figure S3: Clustering metrics and testing performance performance on bursty simulated data, with simulated ground truth cell type used as cluster label. Models trained on 7,200 cells with 800 validation cells for 400 epochs. All metrics were calculated on 2,000 held-out testing cells. **a.** Intra-cluster distance (average over all clusters). **b.** Intra-cluster variance (average over all clusters). **c.** Inter-cluster distance (average over all clusters). **d.** Average inter-cluster distance scaled by average intra-cluster distance. **e.** Average silhouette score (over all cells). **f.** Mean squared error between reconstructed nascent and mature means and observed nascent and mature counts (average over all cells). **g.** Fraction of nearest neighbors (closest cells in latent space) in the same cluster as each cell (average over all cells). **h.** Empirical cumulative distribution function (eCDF) of KL divergence between reconstructed distributions (using *biVI* with bursty likelihood) and simulated ground truth distributions for all 2,000 testing cells, evaluated in individual cell types. **i.** Reconstruction error (negative log likelihood under each model) after 400 training epochs on training data. **j.** Reconstruction error after 400 training epochs on testing data. **k.** Time in seconds to train each model for 400 epochs. **l.** Reconstruction error (negative log likelihood under each model) on train and validation data over training epochs for *biVI* with the bursty likelihood and *scVI* with independent negative binomial likelihoods.

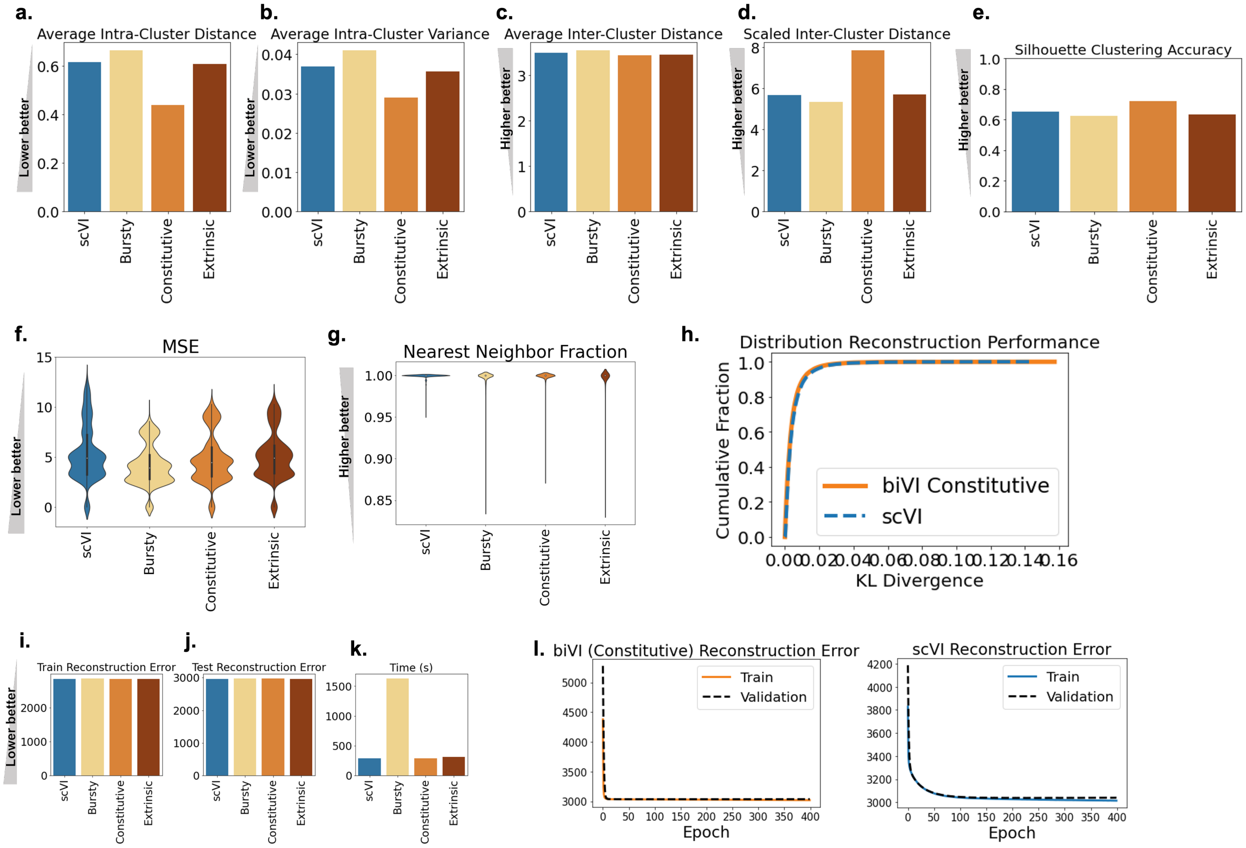


Figure S4: Clustering metrics and testing performance performance on constitutive simulated data, with simulated ground truth cell type used as cluster label. Models trained on 7,200 cells with 800 validation cells for 400 epochs. All metrics were calculated on 2,000 held-out testing cells. All conventions as in Figure S3.

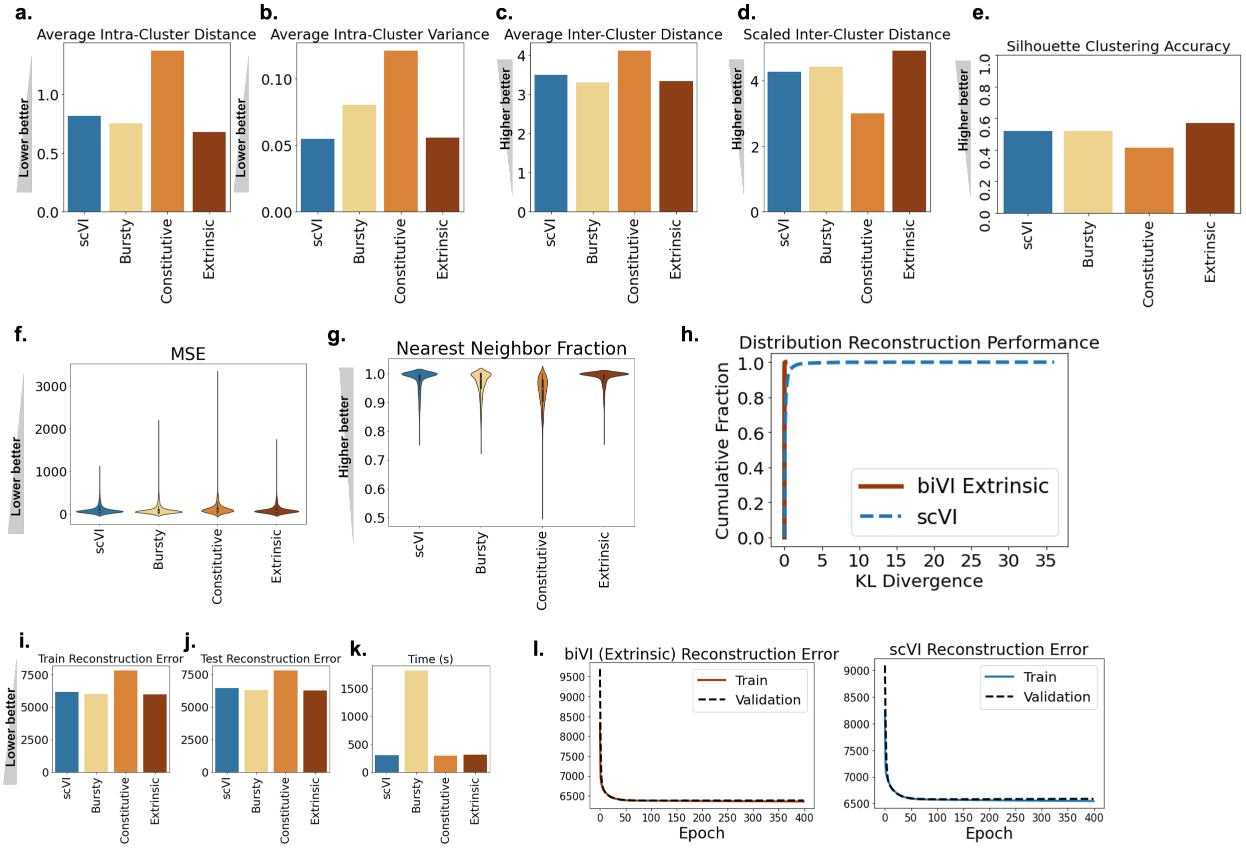


Figure S5: Clustering metrics and testing performance performance on extrinsic simulated data, with simulated ground truth cell type used as cluster label. Models trained on 7,200 cells with 800 validation cells for 400 epochs. All metrics were calculated on 2,000 held-out testing cells. All conventions as in Figure S3.

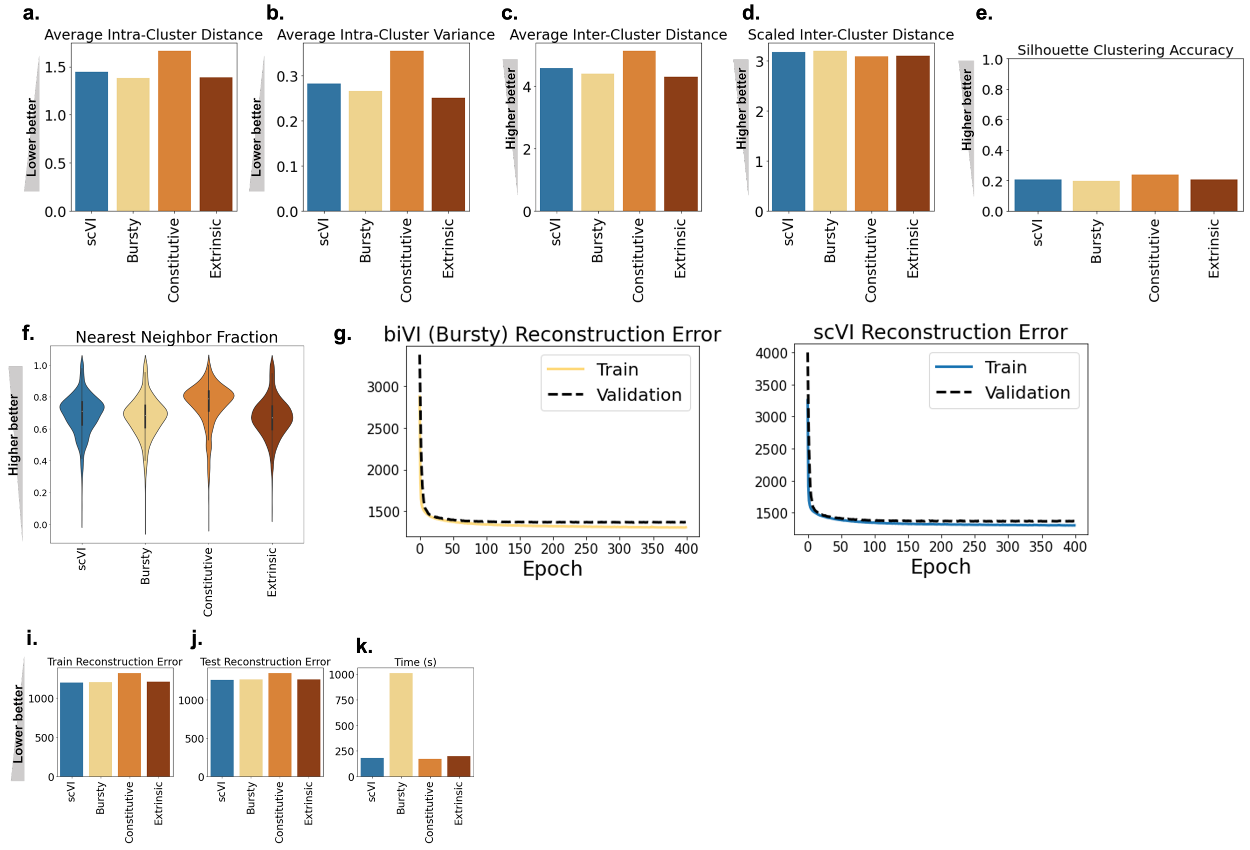


Figure S6: Clustering metrics and testing performance performance on Allen data sample B08, with annotated subclass used as cluster labels. Models trained on 4,622 cells with 513 validation cells for 400 epochs and all metrics calculated on 1,283 held-out testing cells. **a.** Intra-cluster distance (average over all clusters). **b.** Intra-cluster variance (average over all clusters). **c.** Inter-cluster distance (average over all clusters). **d.** Average inter-cluster distance scaled by average intra-cluster distance. **e.** Average silhouette score (over all cells). **f.** Mean squared error between reconstructed nascent and mature means and observed nascent and mature counts (average over all cells). **g.** Fraction of nearest neighbors (closest cells in latent space) in the same cluster as each cell (average over all cells). **h.** Reconstruction error (negative log likelihood under each model) on train and validation data over training epochs for *biVI* with the bursty likelihood distribution and *scVI* with independent negative binomial likelihood distributions. **i.** Reconstruction error (negative log likelihood under each model) after 400 training epochs on training data. **j.** Reconstruction error after 400 training epochs on testing data. **k.** Time in seconds to train each model for 400 epochs.

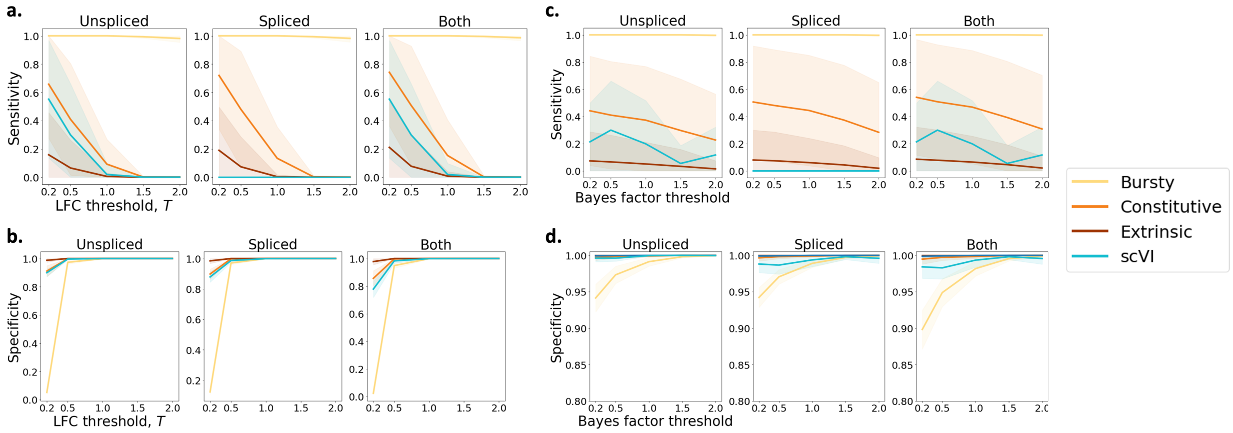


Figure S7: *biVI* with the bursty generative distribution discovers more marker genes for bursty simulated cell types than *biVI* with constitutive and extrinsic generative distributions and *scVI* with independent negative binomial distributions. See Section S6.8 for definitions and discovery procedure. Results shown for genes marked as differentially expressed in inferred unspliced mean, spliced mean, or the union of the two (labeled "both"). **a.** Average sensitivity, or true positive rate, decreases as $log_2$ fold change threshold $T$ increases (with Bayes factor threshold a constant 0.7). Sensitivity was calculated for each cell type: the average over cell types (colored line) plus and minus one standard deviation (lightly shaded) is shown, colored by model (key right of subplots). **b.** The same, but specificity, or true negative rate, which increases as $T$ increases. **c.** Average sensitivity also decreases as Bayes factor threshold increases (with $T$ held constant at 0.5). Again, average and standard deviation over cell types is shown for each model. **d.** Average specificity increases as Bayes factor threshold increases (with $T$ held constant at 0.5).

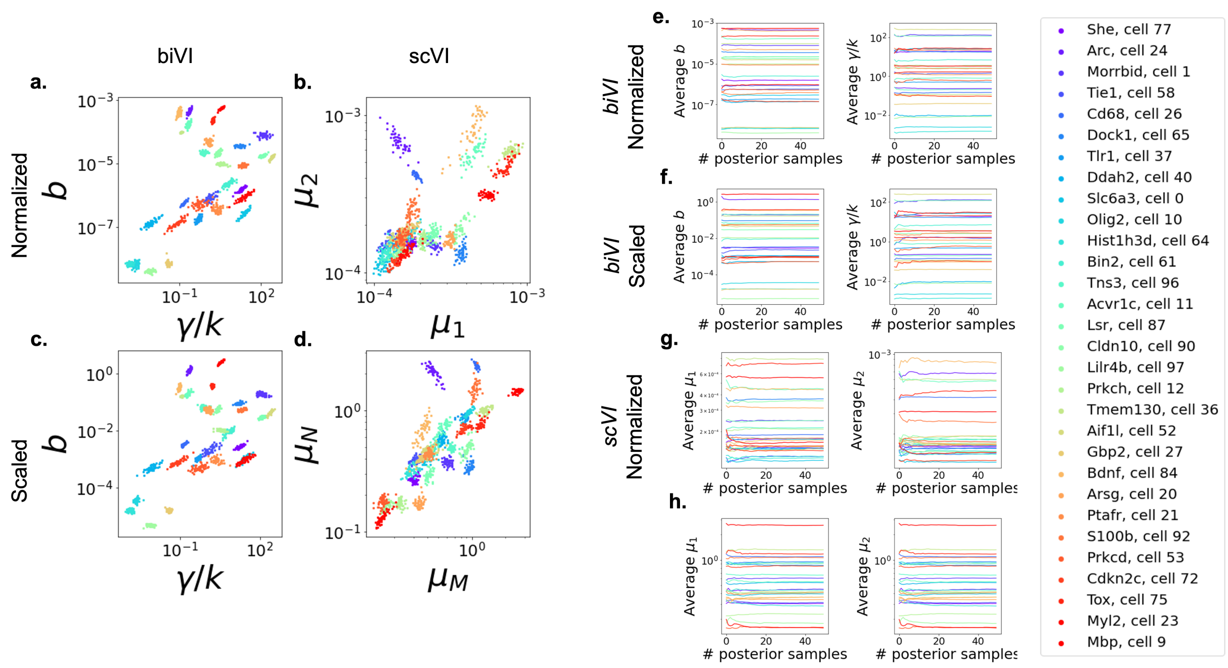


Figure S8: Monte Carlo sampling from posterior of trained *biVI* with bursty generative model and *scVI* model with negative binomial data conditional likelihood does not result in significance variance in inferred conditional parameters. All parameters are for 30 randomly chosen gene-cell pairs and all panels are colored by gene-pair identity (key shown on right). **a.** *biVI* conditional parameters ("normalized", or pre-scaling by sequencing depth). 50 Monte Carlo samples from each cells' posteriors were decoded to produce normalized burst size and relative degradation rates. **b.** *scVI* decoded inferred mean parameters ("normalized", or pre-scaling by sequencing depth). Again, 50 Monte Carlo samples from each cell's posterior were decoded to produce normalized nascent and mature means. **c.** Scaled (post multiplication by sequencing depth) versions of *biVI* inferred parameters. **d.** Scaled (post multiplication by sequencing depth) versions of *scVI* inferred parameters. **e.** Average of *biVI* normalized conditional burst size (left) and relative degradation rate (right) taken over 1-50 posterior samples. Average values converge after several samples. **f.** Average of *scVI* normalized inferred nascent (left) and mature (right) means taken over 1-50 posterior samples. **h.** Average of *biVI* inferred conditional burst size (left) and relative degradation rate (right) taken over 1-50 posterior samples. **i.** Average of *scVI* inferred nascent (left) and mature means (right) taken over 1-50 posterior samples.

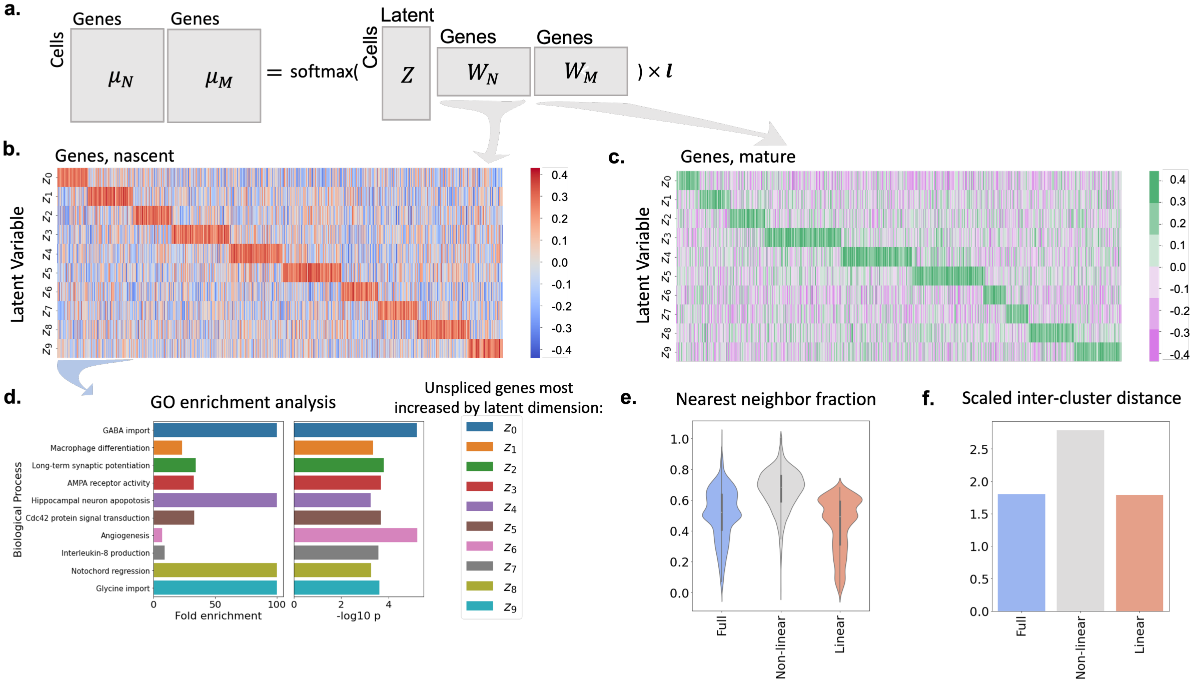


Figure S9: A linear decoder can be used to interpret the effects each latent variable has on gene mean parameters. Discussion of training in Section S9. **a.** Representation of the recovery of means from latent representations with a linear decoder. Sampled latent representations of cells $Z$ are multiplied by weight matrices $W_N$ and $W_M$ and multiplied by sampled log-sequencing depth $l$ after a softmax is applied to produce nascent means $\mu_N$ and $\mu_M$. **b.** Heatmap of weights of linear decoder for nascent means. Genes grouped according to which latent variable ($z_0$ to $z_9$) most greatly increases it (which latent variable is multiplied by highest weight in decoder for nascent mean). **c.** Weights of linear decoder for mature means. Genes again grouped according to which latent variable ($z_0$ to $z_9$) most greatly increases it. **d.** Gene ontology enrichment analysis [79] was performed for modules of genes grouped according to which latent variable most greatly increased their nascent mean. Fold enrichment and negative $log_{10}$ of $P$-value is plotted for the top discovered biological process for each group of genes. **e.** Fraction of nearest neighbors in the same cell type as each cell using observed expression data (nascent and mature means included, labeled "Full"), 10-dimensional latent space obtained from a *biVI* bursty model with three layer non-linear decoder ("Non-linear"), and 10-dimensional latent space obtained from a *biVI* bursty model with a one layer linear decoder ("Linear"). **f.** Average scaled inter-cluster distance (see Section S7), labeled as in **e.**
